# The efficacy of antimicrobial agents is decreased in a polymicrobial environment

**DOI:** 10.1101/2021.07.20.453069

**Authors:** Thomas James O’Brien, Wendy Figueroa, Martin Welch

## Abstract

The airways of people with cystic fibrosis (CF) often harbour diverse polymicrobial communities. These airway infections can be impossible to resolve though antibiotic intervention, even though isolates of the individual species present are susceptible to the treatment when tested *in vitro*. This suggests that susceptibility to antimicrobial agents may be altered in the presence of other microbial species. In this work, we investigate how polymicrobial cultures of key CF-associated species respond to challenge with species-specific antimicrobial agents; colistin (targets *Pseudomonas aeruginosa*), fusidic acid (targets *Staphylococcus aureus*) and fluconazole (targets *Candida albicans*). We found that, compared with growth in axenic cultures, the target organism was protected (sometimes by several orders of magnitude) from the effect(s) of the antimicrobial agent when grown in a polymicrobial culture. This decreased antimicrobial efficacy in polymicrobial cultures was found to have both phenotypic and inherited components. Whole genome sequencing of the colistin-resistant *P. aeruginosa* isolates revealed single nucleotide polymorphisms and indels in genes encoding lipopolysaccharide (LPS) biosynthesis or pilus biogenesis. Colistin resistance associated with loss-of-function mutations in the LPS biosynthetic gene, *wzy*, could be complemented by expression of the wild-type *wzy* gene *in trans*. Our findings indicate that the polymicrobial nature of the CF airways is likely to have a significant impact on the clinical response to antimicrobial therapy.

## Introduction

Cystic fibrosis (CF) is the most common life-limiting genetic disorder affecting the Caucasian population. The disease is caused by defective activity or targeting of the cystic fibrosis transmembrane conductance regulator (CFTR). Over 1800 mutations in the *CFTR* gene have been linked with CF, although the most commonly-encountered is the ΔF508 mutation (www.genet.sickkids.on.20.ca/cftr/Boyle and De Boeck, 2013, Bell et al., 2020). This gives rise to defective targeting of the CFTR and a subsequent imbalance in chloride transport across the epithelial membrane.

A defective CFTR protein gives rise to a systemic multi-organ disease, although perhaps the most striking manifestation of CF is the overproduction of nutrient-rich mucilaginous secretions in the airways. Coupled with a defective mucociliary escalator, such chronic airway obstructions predispose people with CF to life-long infections. These infections are often polymicrobial and include both bacterial and fungal species (Ahmed et al., 2019, Boutin et al., 2015, Mahboubi et al., 2016, Zhao et al., 2012, Hogan et al., 2016, Jorth et al., 2019, Rogers et al., 2004, Rogers et al., 2010b, Rogers et al., 2010a, Sibley et al., 2006, Zemanick et al., 2017). Airway-associated infections are thought to contribute towards the onset of excessive bouts of immunological activity, termed acute pulmonary exacerbations (APEs) (Stenbit and Flume, 2011). APEs cause significant airway scarring and can lead to a decline in pulmonary function (Conrad et al., 2013, Quinn et al., 2014, Carmody et al., 2013, Carmody et al., 2015, Zhao et al., 2012). As such, the onset of APEs is usually met with aggressive antimicrobial treatment, typically aimed at reducing the burden of the principal pathogen(s) present, such as *Staphylococcus aureus* (SA) in infants and *Pseudomonas aeruginosa* (PA) in adolescents and adults (Stanojevic et al., 2017). However, PA in particular exhibits an exquisite predilection for colonizing the CF airways, and in spite of aggressive clinical interventions, by their late ‘teens, most patients become chronically-colonized with this species. A common observation is that, over time, the polymicrobial diversity of the airways decreases and PA eventually becomes the sole occupant of the niche (Conrad et al., 2013, Lyczak et al., 2002). This decrease in polymicrobial diversity is often accompanied by a worsening of airway function, so somewhat counter-intuitively, maintenance of a high polymicrobial diversity appears to be desirable (Zhao et al., 2012). However, the antibiotic interventions used day-to-day to manage CF-associated airway infections, as well as the more aggressive interventions used to treat APEs, are likely to have an impact on all of the species present, and not just the known pathogens (Filkins and O’Toole, 2015, Lopes et al., 2012, Döring et al., 2004, Hauser et al., 2011, Ciofu et al., 2013, Magalhães et al., 2016).

The development and validation of antimicrobial compounds is usually undertaken with a single pathogen in mind, and the presence of additional species at the site of infection is often disregarded (Committee on New Directions in the Study of Antimicrobial Therapeutics, 2006, Leekha et al., 2011). However, there is increasing evidence that interspecies interactions can modulate the gene expression profile and virulence of pathogens when they are grown in co-culture with other species (Antonic et al., 2013, Armbruster et al., 2016, Baldan et al., 2014, Barnabie and Whiteley, 2015, Beaume et al., 2015, Briaud et al., 2019, Dalton et al., 2011, Diggle et al., 2007, Elias and Banin, 2012, Hotterbeekx et al., 2017, Hibbing et al., 2010, Ibberson et al., 2017, Ibberson and Whiteley, 2020, Korgaonkar et al., 2013, Mastropaolo et al., 2005, O’Brien and Fothergill, 2017, Peters et al., 2012, Weimer et al., 2010). Indeed, even species comprising less than 0.05% (by cell numbers) can significantly alter the production of extracellular molecules by the more abundant species in the mixture (O’Brien and Welch, 2019a). It is also possible that such interspecies interactions may alter the response of pathogens such as PA to antimicrobial agents (Briaud et al., 2019, Leekha et al., 2011, Vega et al., 2013, Fodor et al., 2012, Lopes et al., 2012, Peters et al., 2012, Magalhães et al., 2016).

Effective management of the CF-associated microbiota remains a promising approach for improving the quality-of-life and wellness of people with CF. An better understanding of how growth in a polymicrobial consortium alters the response of pathogens to antimicrobial treatment is therefore essential if we are to develop improved interventions (Renwick et al., 2016). However, the paucity of CF polymicrobial infection models has severely hampered progress in this regard. Even though some microbial species are known to co-habit in the human airways, there is no guarantee that these species can be stably co-cultivated in *ex situ* models (O’Brien and Welch, 2019b). To redress this, we recently developed a simple, continuous-flow model for the *in vitro* co-cultivation of the CF-associated microbiota (O’Brien and Welch, 2019a). Our model enables polymicrobial populations of three distinctly different microbial species (PA; a Gram-negative bacterium, SA; A Gram-positive bacterium, and *Candida albicans*; a dimorphic fungus), to be indefinitely maintained at a stable steady-state, with respect to cell titres, mutation rates and extracellular product accumulation.

In this study we show that the different species in our continuous-flow model are maintained in a metabolically-active state, just as has been observed for PA in CF sputum (Palmer et al., 2005, Palmer et al., 2007, Turner et al., 2015). We also examine the effects of three clinically-relevant, species-specific antimicrobials (colistin, fluconazole and fusidic acid) on steady-state axenic and polymicrobial populations of PA, SA and CA. We find that all three species-specific antimicrobials demonstrate lower activity against their target microorganism in the polymicrobial cultures. The protection afforded by growth in a polymicrobial community appears to have both phenotypic and heritable components. For example, PA mutants displaying essentially complete resistance against the “last resort antibiotic”, colistin, could be readily isolated. Whole genome sequencing of these mutants revealed that they contained SNPs and/or indels in a cluster of genes associated with lipopolysaccharide (LPS) biosynthesis. To further confirm the importance of this gene cluster in conferring colistin resistance, we show that colistin sensitivity can be restored in loss-of-function *wzy* mutants by expression of the wild-type gene *in trans* (*wzy* is an LPS biosynthetic gene). These findings highlight the need to re-assess how antibiotics are applied in polymicrobial infection scenarios, and further reinforce the notion that rare/low abundance species may have a profound effect on the physiology and behaviour of the other co-habitants in the niche.

## Results

### Physiological state of PA in the continuous flow setup

There is evidence to suggest that in the CF airways, the PA population is maintained in an actively-growing state (Rogers et al., 2005). We therefore wondered whether the PA in our continuous flow setup was similarly actively growing, or whether it was maintained in a condition more closely approximating to the stationary phase. One way of determining this is to use quantitative reverse-transcription PCR (RT-PCR) to monitor the relative expression of genes associated with exponential phase- and stationary-phase physiology. We compared the gene expression profile of axenically-grown PA in the continuous-flow setup and in a parallel batch culture (comprising the same vessel and stirring rate, but with no input of fresh medium). The latter was included as a control, because we have previously demonstrated that PA grown in ASM under batch culture conditions reaches the stationary phase by 24 h (O’Brien and Welch, 2019a). From a previously published transcriptomic dataset (Mikkelsen et al., 2007) four PA genes that are strongly up-regulated during exponential growth (*rplM*, *rpoA*, *rpsM* and *sodB*; **Figure 1A**) and four PA genes that are strongly up-regulated during stationary phase growth (*rmsA*, *rpoS*, *rmf* and *sodM*; **Figure 1B**) were selected. By targeting 8 different genes, we anticipated that any discrepancies in the expression of the selected genes due to differences in the growth media employed by Mikkelsen *et al* (AGSY medium) compared with the current study (ASM medium) should be mitigated, and that the general metabolic state of the culture(s) would be obvious. Samples were harvested from each culture system at the indicated times following inoculation and RNA was extracted from each sample for subsequent RT-PCR analysis. By 24 h, the continuous flow culture had reached a stable carrying capacity, whereas the batch culture had entered the stationary phase (**Figure S1**). However, by 24 h, the expression of all genes associated with exponential phase growth was high in the continuous flow cultures, whereas the expression of genes associated with the stationary phase was high in the batch cultures. [The relative expression of ribosomal modulation factor (encoded by *rmf*), which converts the 70S subunit into an inactive 100S dimer during the stationary phase, displayed a more gradual decrease over time, but nevertheless, has a negative relative expression value by the 72 h sampling point.] Taken together, these results suggest that at carrying capacity, the microbial population in the continuous-flow culture vessel is maintained in the exponential phase of growth.

**Figure 1.**
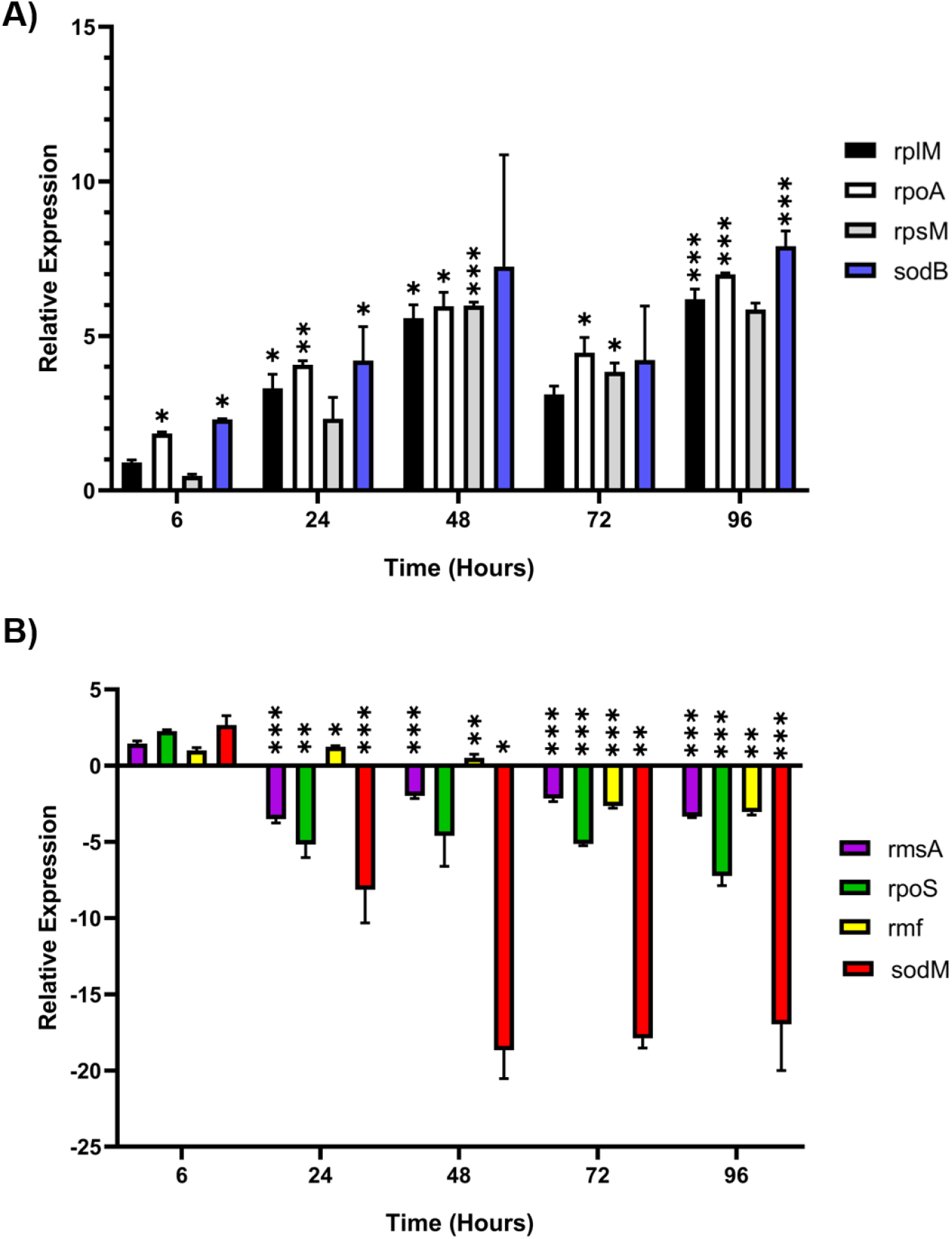
Relative expression of genes associated with exponential and stationary phase growth in PA. The figure shows the fold change expression of the indicated genes in *P. aeruginosa* PAO1 mono-species cultures grown in continuous-flow conditions (*Q* = 170 μL min^−1^) compared with stirred batch conditions (*Q* = 0 μL min^−1^). **(A)** Genes associated with exponential growth; **(B)** genes associated with the stationary phase. Cycle threshold (C_t_) values obtained by quantitative real-time PCR were normalised against expression of the 16S rRNA housekeeping gene and analysed using the comparative ΔΔC_t_ method (Giulietti et al., 2001). Positive values indicate greater expression in the continuous-flow culture; negative values indicate greater expression in the batch culture. Data are the mean ± standard deviation of three independent experiments. Asterix represent significant differences in relative gene expression between cultures maintained under the different growth conditions (* *P* < 0.05, ** *P* < 0.005, *** *P* < 0.001).

### Steady-state microbial communities are attained regardless of microbial inoculation density

The establishment of chronic PA infection is correlated with a worsened patient prognosis and a reduction in the diversity of the microbiota associated with the CF airways (Cox et al., 2010, Zhao et al., 2012). We therefore tested how the introduction of PA at different inoculum densities (OD_600_ 0.05, 0.1, 0.25 and 0.5) perturbs the microbial titres of pre-established SA-CA communities (cultured to a steady-state for 24 h prior to inoculation with PA). There was no appreciable change (P > 0.9) in SA or CA CFU mL^−1^ counts following the introduction of PA to the culture vessel at the lowest inoculation densities (OD_600_ 0.05 and 0.1, **Figure 2A** and **2B** respectively). After 24 h incubation a steady-state microbial population of all three species, with respect to microbial titres, was attained and there was no significant difference (P > 0.9) in PA CFU mL^−1^ at the later points of sampling.

**Figure 2.**
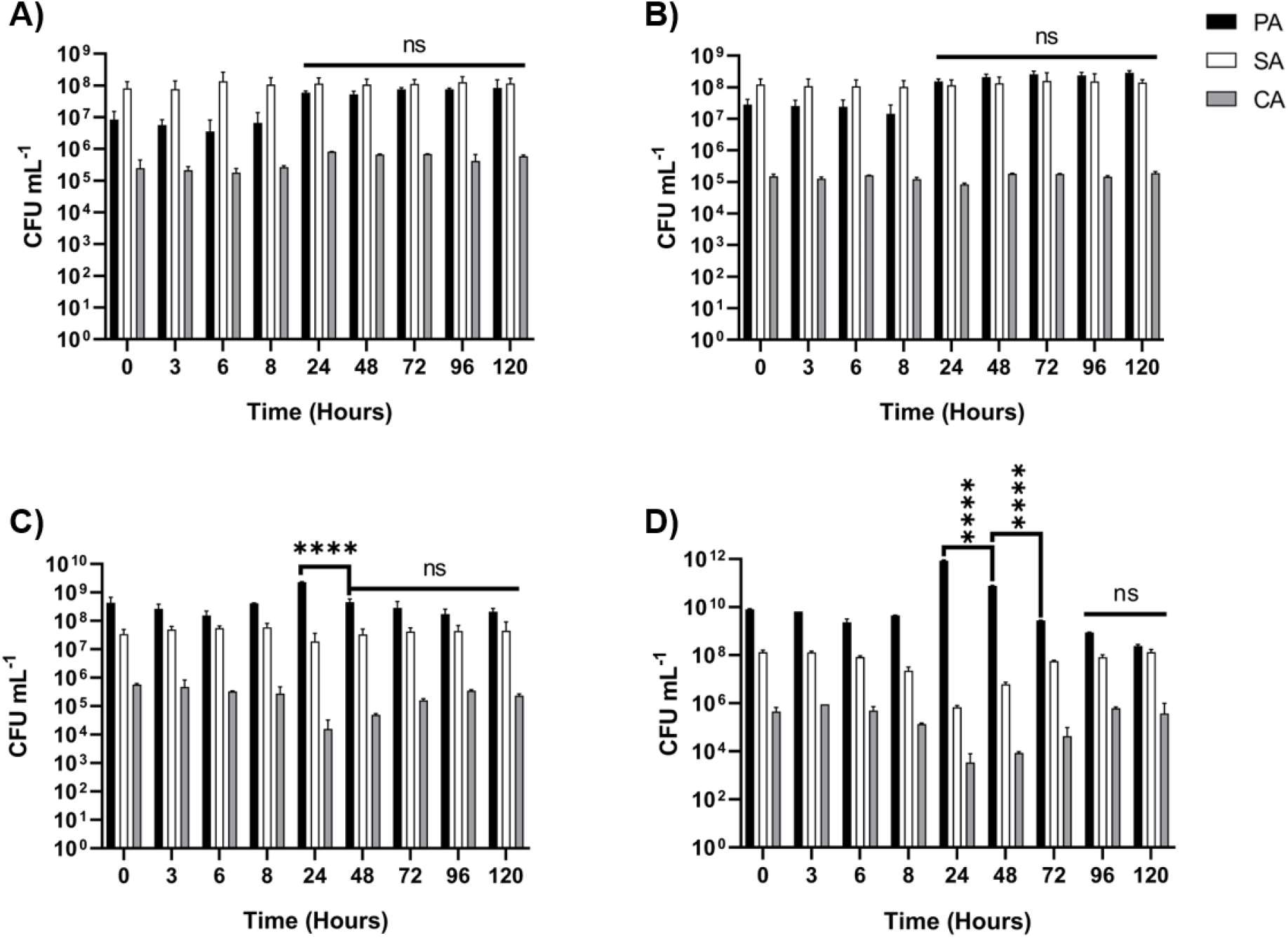
Viable cell counts of individual species following the addition of PA to a steady-state SA-CA co-culture. Dual-species co-cultures of *S. aureus* 25923 and *C. albicans* SC5314 were grown under continuous-flow conditions for 24 hrs. At T = 0 *P. aeruginosa* PAO1 was inoculated into the culture vessel at OD_600_ of 0.05 **(A),** 0.1 **(B)**, 0.25 **(C)** or 0.5 **(D)**. Data represented as mean viable cell counts of PA (black), SA (white) and CA (grey) ± standard deviation following inoculation of PA from three independent experiments. Asterisks represent significant (**** *P* < 0.0001) differences in PA CFU mL^−1^ counts after T = 24 h and in comparison with the previous timepoint. *P* > 0.05 is considered not significant (ns).

As expected, inoculating PA into the culture vessel at the higher densities resulted in significantly higher (P < 0.0001) PA CFU mL^−1^ counts at T = 24 h compared with SA and CA viable cell counts. Inoculation of PA into the culture vessel at an optical density of 0.25 (**Figure 2C**) initially resulted in a 5-fold decrease in SA and CA CFU mL^−1^ counts between T = 0 and 24 h, although this change was not statistically significant (P > 0.9). However, between T = 24 h and 48 h, PA titres began to decline (P < 0.0001) and this was coupled with a recovery in SA/CA CFU mL^−1^ counts. Following the 48 h timepoint an apparently stable steady-state microbial population was established and there was no statistically significant (P > 0.9) change in CFU mL^−1^ counts of any species. Introduction of PA to the steady-state community at OD_600_ 0.5 elicited a similar pattern of resilience. Following addition of the PA, there was initially a significant (P < 0.0001) 10^2^-fold decrease in SA and CA CFU mL^−1^ counts between T = 0 and 24 h (**Figure 2D**). However, this was followed by an approximate 10-fold increase in viable CA and SA CFU mL^−1^ and a significant (P < 0.0001) 10^2^-fold decrease in PA CFU mL^−1^ counts every 24 h after this until T = 96 h. Following this, there was no significant change (P > 0.9) in the CFU mL^−1^ counts of any species. Importantly, there was no significant difference (P > 0.9) in endpoint PA CFU mL^−1^ counts between the co-cultures inoculated with the different densities of PA, demonstrating that steady-state microbial communities, with respect to cell titres, are attained using our continuous-flow model regardless of the initial density of microbial inoculation.

### Antimicrobial minimum inhibitory concentrations in artificial sputum medium

Consistent with our previous study (O’Brien and Welch, 2019a), axenic cultures of *Pseudomonas aeruginosa* PAO1 (PA), *Staphylococcus aureus* ATCC 25923 (SA) or *Candida albicans* SC5314 (CA) and polymicrobial co-cultures of all 3 species reached a steady-state with respect to viable cell counts by 24 h incubation (Figure S2A). There was no significant difference (P > 0.9) in CFU mL-1 counts of any species following the 24 h point of sampling. We next determined the minimum inhibitory concentration (MIC) of three species-specific antimicrobial compounds against PA, SA and CA grown artificial sputum medium (ASM) (Table 1). These data confirmed that colistin (MICPA = 4 μg mL^−1^) has little activity against SA or CA, fluconazole (MIC_CA_ = 1 μg mL^−1^) has little activity against PA or SA, and that fusidic acid (MIC_SA_ = 15.6 ng mL^−1^) has little activity against PA or CA. These MIC values are identical to the clinical breakpoints reported by the European Centre for Antimicrobial Susceptibility Testing using cation-adjusted Müller-Hinton II broth as the culture medium (EUCAST, 2020).

**Table 1.** Minimum inhibitory concentration of antimicrobial compounds. Minimum inhibitory concentration (MIC, μg mL^−1^) of colistin, fusidic acid and fluconazole against *P. aeruginosa* PAO1, *S. aureus* 25923 and *C. albicans* SC5314 grown in ASM. MICs were determined using the EUCAST broth microdilution method. The MIC was considered as the lowest concentration of the compound able to inhibit visible microbial growth after 16 h incubation. MIC values were determined using broth microdilution from three independent experiments using different batches of freshly-prepared ASM each time.

| Species | Minimum Inhibitory Concentration (µg mL<sup>-1</sup>) |  |  |
| --- | --- | --- | --- |
|  | Colistin | Fusidic Acid | Fluconazole |
| *P. aeruginosa* PAO1 | 4 | >256 | >256 |
| *S. aureus* 25923 | >256 | 0.0156 | >256 |
| *C. albicans* SC5314 | >256 | >256 | 1 |

We next compared how single- and mixed-species steady-state microbial cultures responded to treatment with each antimicrobial agent at 2 × and 5 × the determined MIC. [We also tested the effect of adding 1 × MIC of each agent to the cultures (**Figure S3** to **S5**) but this had little impact on cell titres, presumably due to the washout that is an integral feature of the continuous-flow setup.] All microbial populations were grown to a steady-state for 24 h prior to challenge with each antimicrobial agent, which was added by injection through the sterile septum at the top of each culture vessel. It is also important to note that addition of the solvents for each compound were independently confirmed to have no effect on cell titres (**Figure S2B and 2C**). Following addition of the antimicrobial agent, stirring was maintained but the flow was turned off for 1 h to allow the compound time to act before dilution with fresh medium. After 1 h, the flow of fresh medium was restored at the original rate. As fresh ASM enters the culture vessel it gradually dilutes the antimicrobial agent (dilution rate, *D* = 8.7 × 10^−2^ and 0.102 h^−1^ for *Q* = 145 and 170 μL min^−1^, respectively), loosely mimicking the metabolism and excretion of antimicrobials when provided to a patient *in situ*.

### Treatment with colistin

Viable cell counts of pre-established PA mono-species and triple-species populations treated with 2 × MIC (8 μg mL^−1^) or 5 × MIC (20 μg mL^−1^) colistin are shown in **Figure 3**. Following 2 × MIC colistin addition to the mono-species culture, PA titres declined 104-fold over the first 3 h. After this, PA titres increased, and by 24 h - 48 h post-colistin treatment had even reached significantly higher values than those immediately prior to perturbation (**Figure 3A**). A similar sharp decline in PA titres was observed in the mono-species culture treated with 5 × MIC colistin, although here, by 3 h - 5 h post-treatment, no detectable viable PA cells could be recovered from the culture vessels and fewer than 3 PA CFU mL^−1^ were recovered from the culture vessels at 8 h of incubation. However, the population rapidly recovered over the next 16 h. Indeed, by 48 h post-treatment, PA titres were once again significantly higher (P < 0.0001) than the steady-state viable cell counts prior to the addition of colistin (**Figure 3B**).

**Figure 3.**
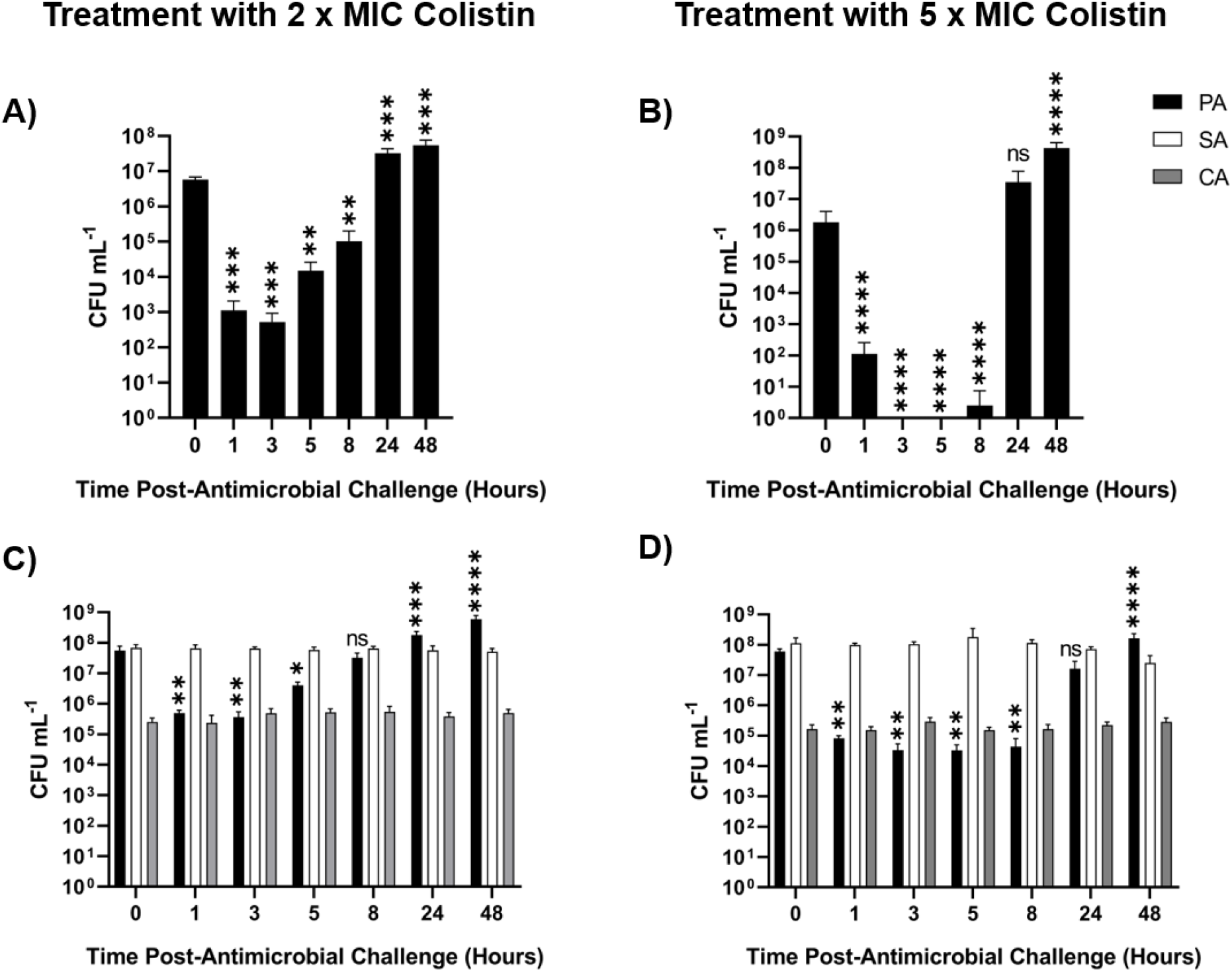
Perturbation of steady-state microbial cultures with 2 x and 5 x MIC colistin. Mono-species *P. aeruginosa* PAO1 (PA) and mixed-species cultures of *P. aeruginosa* PAO1 (black bars), *S. aureus* 25923 (white bars) and *C. albicans* SC5314 (grey bars) were grown to a steady-state in ASM under continuous-flow conditions (24 h incubation). Bars represent viable cell counts (CFU mL^−1^) in **(A, B)** mono-species PA and **(C, D)** mixed-species PA/SA/CA populations following the addition of 8 μg mL^−1^ (2 × MIC) or 20 μg mL^−1^ (5 × MIC) colistin, as indicated, to the culture vessel at T = 0 h. Data represented as the mean ± standard deviation from three independent experiments. Asterisks represent significant differences in PA CFU mL^−1^ counts in comparison to counts at the T = 0 h time point (* *P* < 0.05, ** *P* < 0.005, *** *P* < 0.001, **** *P* < 0.0001).

A very different pattern of response was seen in the colistin-treated polymicrobial cultures. Here, treatment with 2 × MIC colistin led to a 10^2^-fold decrease in PA titres over the first 3 h (i.e., a much smaller decrease than seen in the mono-species culture). Following this, PA titres gradually increased, eventually becoming significantly higher than the starting (pre-treatment) titre. Control cultures, treated with an equal volume of water (used to dissolve the colistin) showed no change in PA/SA/CA titres following addition of the solvent (**Figure S2B**). The impact of colistin was only marginally greater when the cultures were challenged with 5 × MIC; following an initial 10^2^-10^3^-fold decrease in PA titres, the population had completely recovered by 24 h. We conclude that growth in a polymicrobial environment provides protection to PA against the bactericidal effects of colistin. Intriguingly, we saw no significant change (P > 0.9) in SA or CA titres at any point of sampling following challenge with either 2 × MIC or 5 × MIC colistin. This suggests that when the niche normally occupied by PA in the polyculture becomes vacated (due to the action of colistin), this does not necessarily lead to an altered carrying capacity in the other species present.

### Treatment with fusidic acid

Mono-species and triple-species cultures containing SA were challenged with 2 × MIC (31.2 ng mL^−1^) and 5 × MIC (78 ng mL^−1^) fusidic acid (**Figure 4**). Interestingly, there was no change in SA titres in the mono-species culture following treatment with 2 × MIC fusidic acid. However, the effect of fusidic acid on the mono-culture was more pronounced when applied at 5 × MIC, with a ca. 10^2^-fold decrease in SA titres apparent between T = 0 h and 8 h. SA titres then recovered, and by 48 h post-treatment had even significantly surpassed the pre-treatment level by around 5-fold (P < 0.0001).

**Figure 4.**
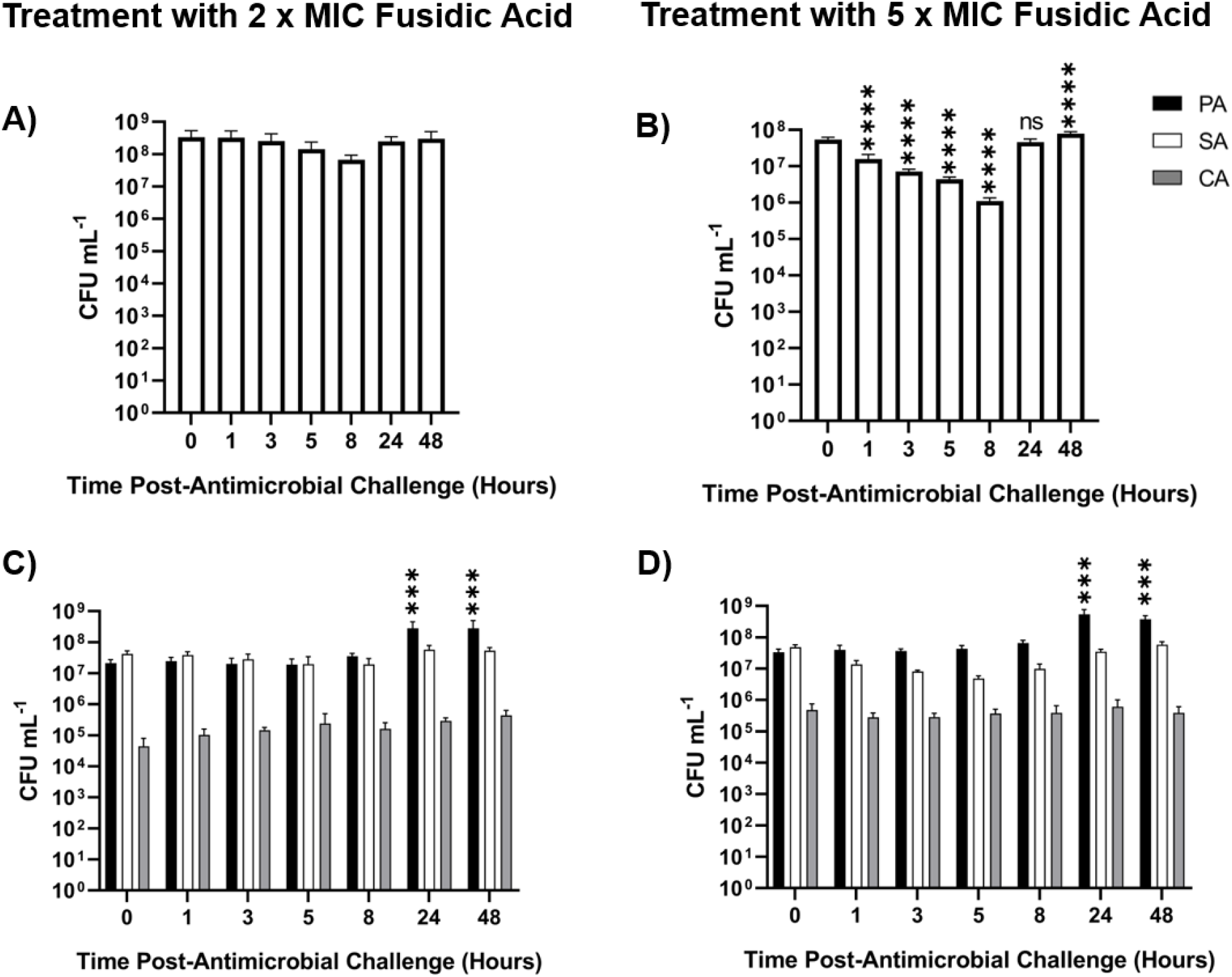
Perturbation of steady-state microbial cultures with 2 x and 5 x MIC fusidic acid. Mono-species *S. aureus* 25923 (SA) and mixed-species cultures of *P. aeruginosa* PAO1 (black bars), *S. aureus* 25923 (white bars) and *C. albicans* SC5314 (grey bars) were grown to a steady-state in ASM under continuous-flow conditions (24 h incubation). Bars represent viable cell counts (CFU mL^−1^) in **(A, B)** mono-species SA and in **(C, D)** mixed-species populations following the addition of 31.2 ng mL^−1^ (2 × MIC) or 78 ng mL^−1^ (5 × MIC) fusidic acid (as indicated) at T = 0 h. Data represent the mean ± standard deviation from three independent experiments. Asterisks represent significant (*** P < 0.001, **** P < 0.0001) differences in SA CFU mL^−1^ counts in comparison to counts at the T = 0 h time point. P < 0.05 is considered not significant (ns).

The situation was different in the polymicrobial cultures. Here, there was no statistically significant change (P > 0.8) in SA or CA titres across any time points for the polymicrobial culture treated with 2 × MIC or 5 × MIC fusidic acid. However, we did observe a significant (P < 0.001) ca. 10-fold increase in PA titres following challenge with 2 × MIC or 5 × MIC fusidic acid when comparing the T = 0 h and T = 24 h samples. [As outlined earlier, control cultures treated with an equal volume of ethanol (used to dissolve the fusidic acid) showed no change in PA/SA/CA titres following addition of the solvent (**Figure S2C**).] This observation suggests that even minor perturbations in the composition or physiology of the non-PA microbial community – in this case, driven by the addition of fusidic acid - may be sufficient to trigger a change in PA titres.

### Treatment with fluconazole

CFU mL^−1^ counts of pre-established CA mono-species and polymicrobial populations treated with 2 × MIC (2 μg mL^−1^) and 5 × MIC (5 μg mL^−1^) fluconazole are shown in **Figure 5**. There was a significant (P < 0.01) ca. 10-fold decrease in CA cell titres recovered from the axenic population treated with 2 × MIC fluconazole following the first 8 h of fluconazole treatment, although titres recovered to pre-treatment levels by the 48 h time point. There was no significant change in CA titres in the polymicrobial population treated with 2 × MIC fluconazole.

**Figure 5.**
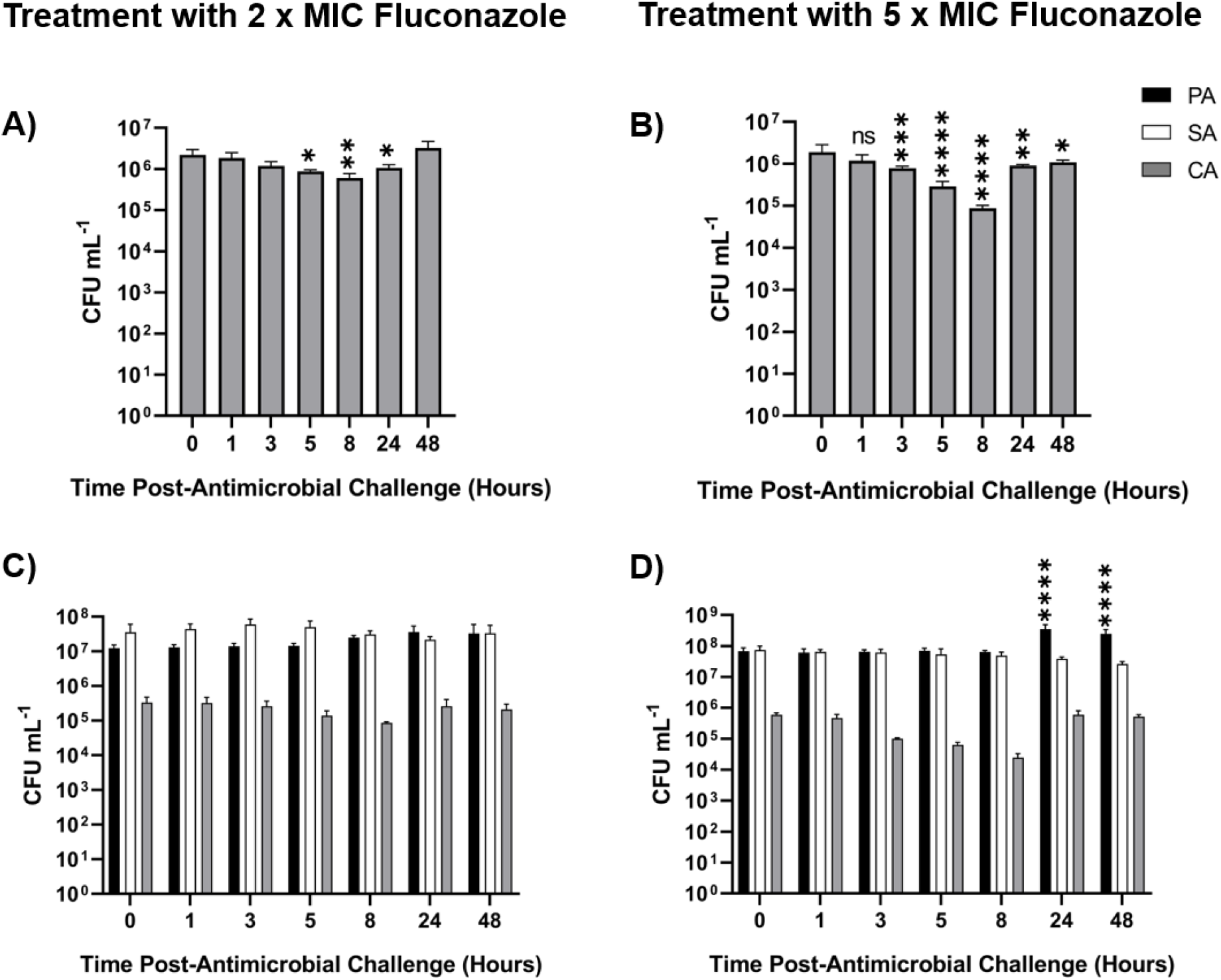
Perturbation of steady-state microbial cultures with 2 x and 5 x MIC fluconazole. Mono-species *C. albicans* SC5314 (CA) and mixed-species cultures of *P. aeruginosa* PAO1 (black bars), *S. aureus* 25923 (white bars) and *C. albicans* SC5314 (grey bars) were grown to a steady-state in ASM under continuous-flow conditions (24 hrs incubation). Bars represent viable cell counts (CFU mL^−1^) within **(A, B)** mono-species CA and **(C, D)** mixed-species populations following the addition of **(A, C)** 2 μg mL^−1^ or **(B, D)** 5 μg mL^−1^ fluconazole to the culture vessel at T = 0 h. Data represented as the mean ± standard deviation from three independent experiments. Asterisks represent significant (* *P* < 0.05, ** *P* < 0.01, *** *P* < 0.001, **** *P* < 0.0001) differences in CA CFU mL^−1^ counts in comparison to counts at the 0 h time point. *P* > 0.05 is considered not significant (ns).

Treatment of the axenic CA culture with 5 × MIC fluconazole led to a more pronounced effect, with a 10^2^-fold decrease in CA titres 8 h after addition of the compound. Titres did recover over the next 40 h, although they remained significantly lower (P < 0.05) than the pre-treatment levels. Challenge of the polymicrobial culture with 5 × MIC fluconazole resulted in a decrease in CA titres, although this decrease was not significant (P > 0.05). However, and similar to the situation seen following challenge of the polymicrobial cultures with fusidic acid, we observed a significant (P < 0.001) ca. 10-fold increase in PA titres following challenge with 5 × MIC fluconazole when comparing the T = 0 h and T = 24 h samples. [Control cultures treated with an equal volume of ethanol (used to dissolve the fluconazole) showed no change in PA/SA/CA titres following addition of the solvent (**Figure S2C**).] This observation reinforces the notion that minor perturbations in the composition or physiology of the non-PA microbial community can trigger a change in PA titres.

Taken together, our findings demonstrate that growth in a polymicrobial consortium confers a high level of protection in sensitive species against the action of microbicidal (colistin) and microbiostatic (fusidic acid, fluconazole) agents. We also conclude that antimicrobial compounds that target non-PA species may confer a selective advantage (in terms of cell titres) on the PA present.

### MIC of recovered isolates

We next sought to determine whether the protection against antimicrobial agents conferred by growth in the presence of co-cultured species was heritable. A selection of 48 isolates were randomly selected from agar spreads generated from the axenic and polymicrobial cultures following challenge with each different antimicrobial agent (applied at 1 ×, 2 ×, and 5 × MIC). Following colony purification, the resistance profile of each isolate was measured and compared against the MIC of the parental strain (**Table 1**) used to inoculate the cultures. A summary of the data is shown in **Tables 2** and **3**.

**Table 2.** Growth of recovered *P. aeruginosa* isolates in ASM with colistin. Number of *P. aeruginosa* PAO1 isolates able to grow in ASM containing the indicated concentration of colistin (in μg mL^−1^). Four isolates were collected at each time point (1 h, 8 h, 24 h and 48 h) after challenge of each culture with 1 ×, 2 ×, or 5 × MIC colistin (i.e., 4 x 4 x 3 = 48 isolates). Isolates were randomly selected from agar spreads of appropriately diluted single-species (upper panel) or PA/SA/CA mixed-species (lower panel) steady-state continuous-flow cultures. The ability of each isolate to grow in different concentrations of colistin was then determined. The results were confirmed over three independent experiments using fresh batches of ASM.

| Time (Hours) | MIC ($\mu\text{g mL}^{-1}$) of colistin against isolates collected from mono-species culture challenged with: | | | | | | | | | | | |
| --- | --- | --- | --- | --- | --- | --- | --- | --- | --- | --- | --- | --- |
|  | 1 x MIC colistin |  |  |  | 2 x MIC colistin |  |  |  | 5 x MIC colistin |  |  |  |
| 1 | 4 | 10 | 20 | 20 | 25 | 45 | >1000 | >1000 | 20 | 25 | 45 | 45 |
| 8 | 4 | 10 | 20 | 20 | 20 | 45 | >1000 | >1000 | 20 | 25 | 20 | 20 |
| 24 | 4 | 4 | 4 | 20 | 20 | 20 | 20 | 20 | 20 | 20 | 20 | 20 |
| 48 | 4 | 4 | 4 | 10 | 20 | 20 | 20 | 20 | 20 | 20 | 20 | 20 |
| | MIC ($\mu\text{g mL}^{-1}$) of colistin against isolates collected from polymicrobial co-culture challenged with: | | | | | | | | | | | |
|  | 1 x MIC colistin |  |  |  | 2 x MIC colistin |  |  |  | 5 x MIC colistin |  |  |  |
| 1 | 4 | 4 | 4 | 10 | 45 | 45 | >1000 | >1000 | 20 | 20 | 25 | 45 |
| 8 | 20 | 20 | 20 | 20 | 45 | >1000 | >1000 | >1000 | 25 | 25 | 45 | 45 |
| 24 | 4 | 20 | 20 | 20 | 25 | 45 | >1000 | >1000 | 20 | 20 | 25 | 25 |
| 48 | 4 | 10 | 20 | 20 | 20 | 20 | 20 | 20 | 20 | 20 | 20 | 20 |

**Table 3.** Growth of recovered *C. albicans* isolates in ASM with fluconazole. Number of *C. albicans* SC3541 isolates able to grow in ASM containing the indicated concentration of fluconazole (in μg mL^−1^). Four isolates were collected at each time point (3 h, 8 h, 24 h and 48 h) after challenge of each culture with 1 ×, 2 ×, or 5 × MIC fluconazole (i.e., 4 x 4 x 3 = 48 isolates in all). Isolates were randomly selected from agar spreads of appropriately diluted single-species (upper panel) and PA/SA/CA mixed-species (lower panel) steady-state continuous-flow cultures. The ability of each isolate to grow in different concentrations of fluconazole was then determined. The results were confirmed over three independent experiments using fresh batches of ASM.

| Time (Hours) | MIC ($\mu\text{g mL}^{-1}$) of fluconazole against isolates collected from mono-species culture challenged with: | | | | | | | | | | | |
| --- | --- | --- | --- | --- | --- | --- | --- | --- | --- | --- | --- | --- |
|  | 1 x MIC fluconazole |  |  |  | 2 x MIC fluconazole |  |  |  | 5 x MIC fluconazole |  |  |  |
| 3 | 1 | 1 | 1 | 1 | 2.5 | 2.5 | 2.5 | 10 | 2.5 | 2.5 | 2.5 | 2.5 |
| 8 | 1 | 5 | 2.5 | 2.5 | 5 | 10 | 10 | 10 | 5 | 5 | 10 | 10 |
| 24 | 1 | 1 | 2.5 | 2.5 | 5 | 5 | 10 | 10 | 2.5 | 5 | 5 | 10 |
| 48 | 1 | 1 | 1 | 1 | 2.5 | 2.5 | 2.5 | 2.5 | 1 | 2.5 | 2.5 | 2.5 |
| | MIC ($\mu\text{g mL}^{-1}$) of fluconazole against isolates collected from polymicrobial co-culture challenged with: | | | | | | | | | | | |
|  | 1 x MIC fluconazole |  |  |  | 2 x MIC fluconazole |  |  |  | 5 x MIC fluconazole |  |  |  |
| 3 | 2.5 | 2.5 | 2.5 | 2.5 | 5 | 10 | 10 | 10 | 5 | 5 | 5 | 10 |
| 8 | 2.5 | 5 | 5 | 10 | 5 | 10 | 10 | 10 | 5 | 5 | 10 | 10 |
| 24 | 1 | 2.5 | 5 | 5 | 5 | 5 | 10 | 10 | 2.5 | 5 | 5 | 10 |
| 48 | 1 | 1 | 1 | 1 | 2.5 | 5 | 10 | 10 | 1 | 1 | 1 | 2.5 |

**Table 4.**
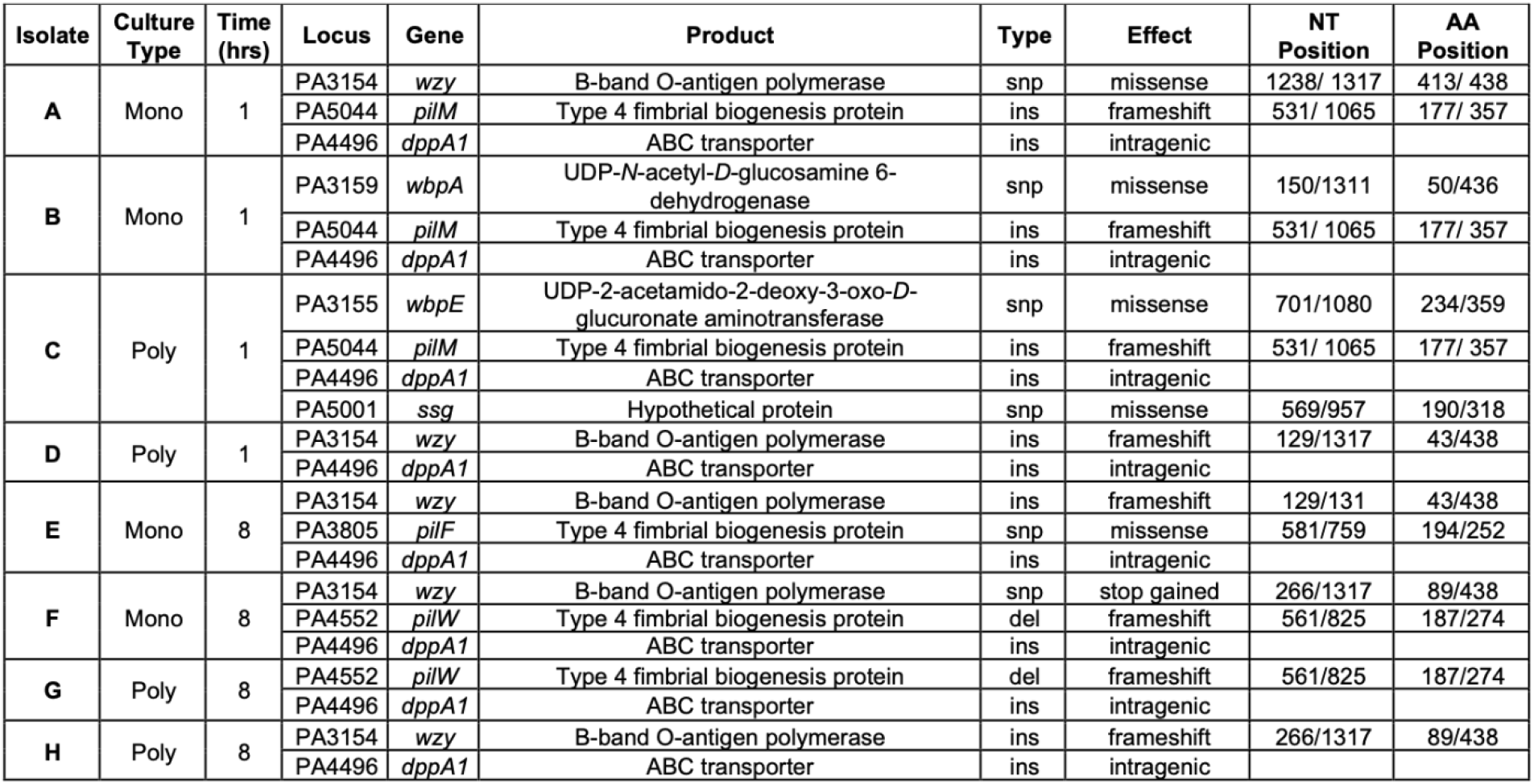
Non-synonymous SNPs and indels associated with the colistin resistant *P. aeruginosa* isolates. Summary of non-synonymous SNPs and indels identified in the genome of PA isolates resistant to the action of 1 mg mL^−1^ colistin when compared with the re-sequenced reference genome of the progenitor (PAO1). All isolates sequenced were collected from a steady-state continuous-flow culture after 1 h or 8 h exposure (as indicated) to 8 μg mL^−1^ colistin. Culture type identifies whether the isolate was obtained from a mono-species (Mono) or polymicrobial (Poly) culture. Mutation type is abbreviated to ins (insertion), del (deletion) or snp (single nucleotide polymorphism). NT position and AA position denote the location of the nucleotide or amino acid change in the gene or encoded protein, respectively.

There was no difference between the MIC of the parental SA strain and any SA isolate collected from either the mono-species or polymicrobial cultures treated with any concentration of fusidic acid (*data not shown*). This suggests that the diminished efficacy of fusidic acid against SA in polymicrobial cultures arises from a protective, not inherited, mechanism of resistance.

This was not the case with the colistin-treated PA. Here, 90% of tested PA isolates from the polymicrobial cultures and 83% of the tested PA isolates from mono-species cultures could grow on the minimum concentration of colistin required to inhibit growth of the parental strain (i.e., 4 μg mL^−1^ colistin, **Table 2**). Indeed, and remarkably, 15% of the PA isolates from the polymicrobial cultures and 8% of the PA isolates from the axenic cultures could grow in ASM supplemented with 1 mg mL^−1^ colistin (i.e., 250 × MIC); we consider these isolates to be fully resistant to colistin. Perhaps even more strikingly, we observed that the prevalence of these fully-resistant mutants was highest in the cultures challenged with 2 × MIC colistin, indicating that the lower concentration of the drug confers more potent selection for resistance than the higher (5 × MIC) concentration. These resistant mutants were identified in the samples harvested 1, 8 and 24 h after colistin challenge, but had disappeared by the 48 h sampling point, perhaps indicating that they are associated with a fitness defect in the absence of selection pressure from the drug. As an aside, we note that we have previously demonstrated that the PA (and SA) in the triple-species steady-state community do not display elevated mutation rates (O’Brien and Welch, 2019a). This suggests that the increased prevalence of mutants resistant to colistin is most likely mediated through selection alone.

A similar, albeit less pronounced, situation was observed with fluconazole and CA (**Table 3**). Here, although no extremely high-level resistance was observed (i.e., an MIC > 10 ◻g mL^−1^), 31% of the tested CA isolates from the polymicrobial cultures and 19% of the tested isolates from the mono-species cultures were able to grow in the presence of 5 μg mL^−1^ fluconazole (cf. MIC_progenitor_ = 1 μg mL^−1^). Once again, we noticed that the highest prevalence of resistance was seen in the cultures treated with the intermediate (2 × MIC) concentration of fluconazole. Taken together, our data suggest that growth as part of a polymicrobial community increases the prevalence of mutants that are resistant to the action(s) of some clinically-relevant antimicrobial agents.

### Whole genome sequencing

The high-level resistance to colistin that we observed was intriguing and in order to identify its possible origins, we used whole genome sequencing (WGS) to identify the genetic changes responsible. Eight PA isolates with MIC > 1mg mL^−1^ were collected from the continuous-flow cultures challenged with 2 × MIC colistin. Two colistin-resistant isolates were collected from the mono-species PA culture after 1 h (isolates A and B) or 8 h (isolates E and F) treatment with colistin, and two resistant isolates were collected from the polymicrobial co-culture after 1 h (isolates C and D) or 8 h (isolates G and H) treatment. As a control, we also re-sequenced an isolate of the progenitor (input) strain of PAO1 that was used to inoculate each culture.

All sequenced isolates contained an insertion (a single cytosine) adjacent to *dppA1* (PA4496), which encodes a dipeptide ABC transporter substrate-binding protein. Although important for the uptake and utilisation of di- and tri-peptides by PA (Pletzer et al., 2014), this insertion occurs immediately after the TAG stop codon of *dppA1* and is therefore predicted to have no effect on the functionality or expression of the gene product. All isolates, except D and H, contain SNPs in one of three genes (*pilM*, *pilW* or *pilF*) associated with the biogenesis of the type IV fimbrial protein (also called type IV pilin). Furthermore, isolates A, D, E, F and H contain SNPs or small indels in *wzy* (PA3154), which encodes for the B-band O-antigen polymerase. Isolates B and C also contained SNPs in the *wzy* cluster; isolate B contained a SNP in *wbpA* (PA3159) and isolate C contained a SNP in *wbpE* (PA3155). Both of these genes are involved in biosynthesis of the O-antigen oligosaccharide. Staying with this theme, isolate C also contained a T→C transition in *ssg* (PA5001), which, although not well-characterized in PA, encodes a putative glycotransferase involved in the biosynthesis of LPS in *Pseudomonas alkylphenolia* (Veeranagouda et al., 2011). Taken together, our data strongly suggest that colistin resistance in both mono-species and polymicrobial cultures is mediated through mutations in the genes associated with LPS or pilus biogenesis.

### Complementation of the colistin-resistant phenotype

In order to test our hypothesis regarding the involvement of the LPS biosynthetic pathway in colistin resistance, we carried out complementation experiments to determine whether the susceptibility to colistin could be restored by expression of a wild-type version of the affected gene. Since most (5 out of 8) of the isolates had mutations in *wzy*, we introduced the wild-type version of this gene into the shuttle vector, pUCP20, and expressed this in isolates D and F. These isolates have a frameshift mutation and a premature stop codon (respectively) early on in the *wzy* ORF, and likely lead to gross loss-of-function of the gene. As expected, the MIC of isolates D and F for colistin was high in the absence of the wild-type *wzy* gene. However, the presence of a functional *wzy* (introduced on pUCP20-*wzy*) led to a 32-fold reduction in the MIC of colistin in both ASM and Mueller-Hinton broth (MHB) compared with the isolates transformed with an empty plasmid (**Figure 6**). This indicates that the loss of function of *wzy* in these isolates is indeed responsible for the observed colistin resistance phenotype. Taken together, these data demonstrate that the LPS of *P. aeruginosa* plays a crucial role in colistin susceptibility and that loss-of-function mutations in the genes involved in LPS biosynthesis represent a novel mechanism of resistance to this antibiotic.

**Figure 6.**
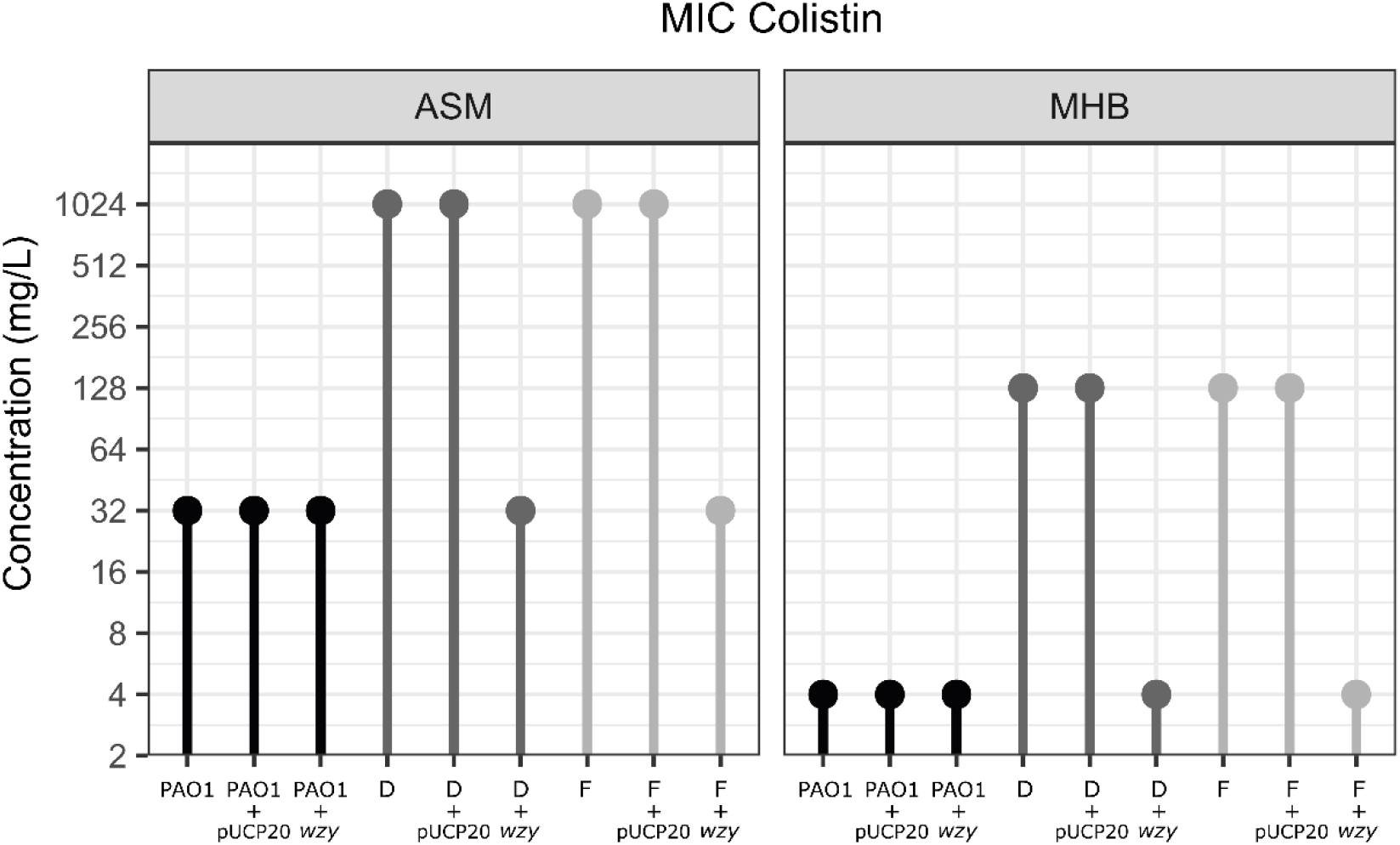
MIC of colistin for colistin-resistant isolates. The minimal inhibitory concentration of colistin is shown for the parental strain (PAO1) and for two of the colistin-resistant isolates (D and F, shown in different shades of grey). The MICs were determined using the microdilution/resazurin method (see methods) for each indicated strain containing either no plasmid, or empty pUCP20, or pUCP20-*wzy*. Experiments were performed using three biological replicates.

## Discussion

To date, very few studies have examined how antimicrobial treatment perturbs mixed microbial communities grown in defined, *in vitro* conditions. This is largely due to the technical difficulties associated with setting up polymicrobial cultures that display long-term stability (O’Brien and Welch, 2019b). Earlier researchers have examined the response of polymicrobial biofilms to therapeutic intervention and report an altered, and often refractory, response to antimicrobial treatment (Sharma et al., 2019). These studies often cite decreased diffusion of compounds through the biofilm matrix (Adam et al., 2002, Kong et al., 2016, Kean et al., 2017), increased horizontal gene transfer of antimicrobial resistance genes (Tanner et al., 2017, Ready et al., 2006, Weigel et al., 2007, Sørensen et al., 2005) and an increased concentration of antimicrobial enzymes, e.g. β-lactamases (Sorg et al., 2016, Perez et al., 2014, Lee et al., 2013), in biofilms as the reason for modulated responses to therapeutic intervention. However, biofilms exhibit heterogenous gene expression profiles (Stewart and Franklin, 2008), and their metabolism is broadly considered to be similar to cultures in the stationary phase of growth (Jaishankar and Srivastava, 2017, Anderl et al., 2003). This may be significant since antimicrobial agents are generally less effective against both biofilms and stationary phase cells (Kamble and Pardesi, 2020, Anderl et al., 2003, Høiby et al., 2010, Leekha et al., 2011, Lopes et al., 2012). Indeed, a recent study of how planktonic and biofilm SA-CA co-cultures respond to antimicrobial treatment also found that growth as a co-culture reduces the susceptibility of SA to antimicrobials that readily diffuse through the biofilm matrix (Nabb et al., 2019). The authors report that a reduced metabolic activity of SA during co- culture (as a result of faster nutrient depletion during co-culture) is the reason for this observed decrease in antimicrobial efficacy. By contrast, in the current study, we demonstrate that steady-state planktonic populations can be maintained in the exponential phase of growth indefinitely when using the continuous-flow setup described by (O’Brien and Welch, 2019a). As previously noted, this is important because the PA population in the CF airways is known to be maintained in an actively-growing state (Rogers et al., 2005). Indeed, to the best of our knowledge, this study is the first to examine the impact of external perturbations such as antimicrobial agents on a stable, yet metabolically-active CF-associated polymicrobial community. Such studies have not been possible using existing *in vitro* or *in vivo* models of CF airway infections.

We demonstrate a clear decrease in the efficacy of three clinically-utilised, species-specific antimicrobial compounds (colistin, fusidic acid and fluconazole) against their target microorganism when that organism is grown as part of a polymicrobial consortium. This observation highlights the need to consider the role of additional species at the site of a polymicrobial infection, and how these might alter the response of a principal pathogen to a previously validated therapeutic intervention. Given the diverse polymicrobial communities associated with CF-airway infections, this finding can, in part, explain the failings of current antimicrobial treatment regimens in eradicating PA infections. We also provide evidence that species-specific antimicrobial interventions can have unintended consequences on the dynamics of those species that are not targeted by the drug. These consequences are not necessarily predictable either – niche vacation *per se* is not necessarily accompanied by niche infiltration by the other species present.

Modulations in antimicrobial efficacy in the polymicrobial environment were found to involve protective (phenotypic) and inherited elements of tolerance/resistance. For example, although growth in the polymicrobial culture appeared to confer protection upon SA against fusidic acid, there was no difference in the antimicrobial susceptibility of the parental strain or any recovered isolate. This indicates a protective (non-heritable) mechanism of tolerance that requires the presence of PA and/or CA in the co-culture. By contrast, a high proportion of the PA or CA isolates challenged with colistin or fluconazole (respectively) displayed heritable resistance against these agents.

WGS analysis of PA isolates resistant to the action of 1 mg mL^−1^ colistin revealed SNPs/indels in a set of genes common also to isolates recovered from axenic cultures. This finding demonstrates that growth of PA as part of a polymicrobial consortium does not necessarily drive the emergence of different inherited mechanisms of colistin resistance. However, growth in the polymicrobial culture did increase the frequency of PA isolates that were resistant to colistin (this, in spite of our earlier observations that the PA mutation rate in polymicrobial cultures is the same as that in axenic cultures). Interestingly, in the colistin-resistant isolates that were subjected to WGS, the genes affected by SNPs and indels were all involved in the biogenesis of LPS or pilin. Modification of LPS through the addition of 4-amino-L-arabinose to exposed phosphate groups on the lipid A moiety has been previously implicated in conferring resistance to cationic antimicrobial peptides such as colistin (Olaitan et al., 2014). However, this L-Ara4N modification is associated with a dedicated gene cluster (*arnBCADTEF-pmrE*; Raetz et al., 2007). To our knowledge, the current study is the first to prove that mutations in the core LPS biosynthetic pathway genes can also give rise to high levels of colistin resistance. Given the importance of colistin as a last-resort antibiotic, this highlights the need for more fundamental studies aimed at establishing the exact role of LPS in colistin susceptibility. Additionally, our results underscore the importance of carrying out evolution experiments to assess antimicrobial resistance in settings that better mimic the environmental conditions encountered during infections. SNPs in genes involved in biosynthesis of type IV fimbrial proteins were also found in most of the sequenced PA isolates in this study. Interestingly, mutations in the pilin biogenesis genes have been previously identified in two separate WGS-based studies of colistin-resistant clinical PA isolates (Lee et al., 2014, Lee et al., 2016). However, and beyond this correlation, there is currently no direct evidence linking pilin synthesis with colistin resistance. Given that isolate G in the current study could grow in the presence of 1 mg mL^−1^ colistin, yet only contains a non-synonymous mutation in *pilW*, this provides strong evidence to suggest that disruption of type IV pilin synthesis can also give rise to colistin-resistance. In summary, we have identified novel SNPs and indels in seven genes involved in the synthesis of LPS or pilin (*wzy*, *wbpA*, *wbpE*, *ssg*, *pilF*, *pilM* and *pilW*) that likely contribute towards or account for resistance to colistin.

The LPS and pili biosynthetic genes are known to be important for mediating host-bacterium interactions (Lam et al., 2011), although they are not essential for the growth of PA *in vitro* (Ramphal et al., 1991). Interestingly, mutations in LPS and pilin biosynthetic genes are common among clinical PA isolates recovered from late-stage CF airway infections (Smith et al., 2006). Moreover, PA clinical isolates from late-stage CF airway infections are often defective in virulence factor production, and in batch cultures, can be co-cultivated with other bacterial species for longer periods of time (Baldan et al., 2014). We therefore hypothesise that down-regulation of genes encoding cell surface moieties may benefit PA when it is growing as part of a polymicrobial consortium. Speculatively, this may be by preventing effective recognition of PA by other members of the community. If such recognition is accompanied by up-regulation of interspecies competition mechanisms, it is easy to see why this strategy may be of benefit to PA (Korgaonkar et al., 2013). This may also explain why mutants defective in such genes would be expected to be more prevalent in steady-state polymicrobial co-cultures.

For the populations treated with 5 × MIC fusidic acid there was a significant increase in PA cell titres in the polymicrobial community at the 48 h sampling point (compared with pre-treatment titres). This demonstrates the ability of PA to expand into an environmental niche that has been vacated following antimicrobial agent-mediated perturbation of the polymicrobial community. This key finding supports the climax-and-attack model (CAM) of changes within a polymicrobial community following clinical intervention or immune clearance proposed by Conrad et al. (2013). Briefly, this hypothesis suggests that keystone pathogens of late-stage CF airway infections can resist clearance from the airway and acquire adaptations that enable them to occupy a newly-liberated environmental niche. This often complicates future treatment regimens and leads to a worsening patient prognosis. Given that fusidic acid challenge caused a decrease in SA titres and a concomitant increase in PA titres, this observation further affirms the importance of understanding how the entire polymicrobial population, and not simply a target pathogen, responds to antimicrobial treatment.

Our finding, that the PA genes affected by SNPs and indels in the continuous-flow polymicrobial cultures are similar to those associated with PA isolates from chronic CF infections suggests that the *in vitro* model captures key elements of the airway environment. Although only a small subset of species and antimicrobial perturbations were investigated in the current study, any number of species and compound combinations can, in principle, be tested. One key future application of the setup will be to examine how patient-derived polymicrobial consortia respond to antimicrobial-mediated perturbation, and efforts are currently underway to develop this approach. Additionally, it stands to reason that if the co-culture of some microorganisms can increase a species’ ability to tolerate therapeutic intervention, the inverse trend may also be true. Indeed, by promoting the growth of “beneficial” species in the CF microenvironment, it may be possible to increase the antimicrobial susceptibility of “keystone” respiratory pathogens. Although previously near-impossible to test, this goal becomes more realistic using the *in vitro* reconstitution approach that we describe here.

In summary, we have developed a simple, robust model that enables real-time facile examination of how metabolically-active microbial communities respond to exogenous perturbation. Ultimately this enables the role of co-cultivated microbial species to be explicitly considered when developing/validating novel treatment regimens for the effective management of polymicrobial infection scenarios.

## Materials and Methods

### Microbial strains and growth media

The bacterial/fungal strains used in this study were: *P. aeruginosa* PAO1 (PA; strain MPAO1 from Colin Manoil), *S. aureus* ATCC 25923 (SA) and *C. albicans* SC314 (CA). All strains were routinely cultured in lysogeny broth (Formedium) with vigorous aeration at 37°C overnight. Artificial sputum medium (ASM) was freshly prepared for every experiment as described in (O’Brien and Welch, 2019a).

### Culture conditions

The continuous-flow culture vessel was assembled, inoculated and incubated as described in (O’Brien and Welch, 2019a). Briefly, overnight cultures were washed three times in sterile 1 × phosphate buffered saline (PBS, Oxoid) prior to inoculating the culture vessel. Pre-warmed ASM (100 mL) in the culture vessels was inoculated with the required combination of microbial species. Unless otherwise indicated, each species was introduced into the culture vessel to achieve a starting OD_600_ of 0.05. The vessel was incubated for 3 h prior to staring the flow of medium. For experiments not containing CA, the flow rate (*Q*) was set at 170 μL min^−1^. For experiments including CA, *Q* was decreased to 145 μL min^−1^. Samples (1 mL volume) were withdrawn using a syringe fitted with a sterile needle inserted through the rubber septa in the HPLC ports. Batch cultures were set up as described for the continuous-flow culture experiments, except that *Q* = 0 μL min^−1^. For all experiments, the culture temperature was maintained at 37°C.

### Quantitative real-time PCR (RT-PCR)

Single-species cultures of PA were grown in ASM under continuous-flow and batch culture conditions as described above. Cells were harvested by removing 2 mL of the culture and pelleting *via* centrifugation (13,000 × *g*, 1 min, 4°C). RNA was extracted using an RNeasy Plus Mini Kit (Qiagen) following the manufacturers’ bacterial extraction protocol. cDNA synthesis was performed in a single-step reaction using the High-Capacity cDNA Reverse Transcription Kit (ThermoFisher) following the manufacturers’ instructions. To determine the physiological state (actively growing or stationary phase) of the harvested cells, four PA genes whose expression is associated with stationary phase growth and four PA genes whose expression is associated with exponential phase growth were selected from a previously published transcriptomic dataset (Mikkelsen et al., 2007).*Supplementary information 6* details the target genes, primers and quantitative real-time PCR (RT-PCR) reaction conditions used. Cycle threshold (C_t_) values of cDNA samples obtained by RT-PCR were analysed using the comparative ΔΔC_t_ method (Giulietti et al., 2001). For each target gene, C_t_ values were normalised according to the C_t_ value of the constitutively expressed 16S rDNA housekeeping gene encoding for the 30S ribosomal subunit (Clarridge, 2004) on the same reaction plate.

### Introduction of PA to steady-state SA-CA co-cultures

Dual species co-cultures of *S. aureus* 25923 and *C. albicans* SC5314 were grown to a steady-state for 24 hrs under continuous-flow conditions (as described above). Routine overnight cultures of *P. aeruginosa* PAO1 were then washed three times in PBS and introduced into the culture vessel to final OD_600 nm_ 0.05, 0.1, 0.25 or 0.5. The culture vessel was then incubated under continuous-flow conditions, *Q* = 145 μL min^−1^.

### Determination of antimicrobial minimum inhibitory concentrations (MICs)

The minimum inhibitory concentration (MIC) of colistin, fluconazole and fusidic acid was determined in ASM using the broth microdilution method described by the Clinical and Laboratory Standards Institute (CLSI, 2019). Briefly, stock solutions of the antimicrobial agents were freshly prepared prior to each experiment. Serial 2-fold dilutions (in fresh, pre-warmed ASM) of each antimicrobial agent were then made and dispensed into a 96-well microtiter plate (Nunc). Overnight cultures of each microbial strain were washed three times in sterile PBS and used to inoculate the wells to an initial OD_600_ of 0.05. The final volume of liquid in each well was 150 μL. The plates were sealed with a gas permeable Breathe-Easy membrane (Sigma) and incubated for 16 h at 37°C with 100 rpm shaking. The MIC was taken as the lowest antimicrobial concentration able to inhibit visible microbial growth.

### CFU mL^−1^ enumeration

Colony forming units (CFU) per mL of culture were determined using the single plate-serial dilution spotting (SP-SDS) method (Thomas et al., 2015) onto the same selective agar media described in (O’Brien and Welch, 2019a).

### Perturbation of steady-state communities

Mono-species or triple-species cultures of *P. aeruginosa* PAO1, *S. aureus* 25923 and *C. albicans* SC5314 were grown until they reached a steady-state (24 h post-inoculation) under continuous-flow conditions (as described above). Pre-established steady-state microbial cultures were then treated with the appropriate antimicrobial compound at a final concentration of 1 ×, 2 × or 5 × MIC. The antimicrobial agent was introduced through a sterile needle inserted through the rubber septum in the HPLC port. Cultures were then incubated for 1 h with no flow (*Q* = 0 μL min^−1^). An appropriate flowrate, Q = 170 μL min^−1^ for PA and SA mono-species cultures and Q = 145 μL min^−1^ for CA mono-species and triple-species cultures, was then applied to the culture vessel and unless otherwise indicated, the setup was incubated for 48 h. Stock concentrations of colistin (10 mg mL^−1^ in ddH2O, Sigma), fluconazole (10 mg mL^−1^ in ethanol, Sigma) and fusidic acid (10 mg mL^−1^ in ethanol, Sigma) were made fresh prior to each experiment.

### Whole genome sequencing (WGS)

Eight PA isolates able to grow in the presence of 1 mg mL^−1^ colistin were randomly selected for whole genome sequencing (WGS). As a control, we also carried out WGS on the parental strain, PAO1. Two resistant isolates were selected from: the mono-species culture after 1 h exposure to 8 μg mL^−1^ colistin (isolates A and B); the mono-species culture after 8 h exposure to colistin (isolates E and F); the polymicrobial co-culture after 1 h exposure to colistin (isolates C and D) and the polymicrobial co-culture after 8 h exposure to colistin (isolates G and H). WGS of the isolates was performed by MicrobesNG (Birmingham, UK) using the Nextera XT library prep protocol v.05 (Illumina) on an Illumina MiSeq platform using 2 x 250 bp paired-end reads. The reads were adapter-trimmed using Trimmomatic v0.30 with a sliding window quality cut-off of Q15. Taxonomic classification of sequences and assessment of sequence contamination was performed using Kraken (Wood and Salzberg, 2014). The *de novo* assembly of contigs was performed using SPAdes v3.14.0 with the default parameter settings and automated annotation of the resulting contigs was performed using Prokka v1.12. Variants were called using Snippy v2.5/Freebayes v0.9.21-7 (https://github.com/tseemann/snippy) with a minimum base quality of 20, read coverage of 10× and a 90% read concordance at a locus for a variant to be reported. Variants in the different isolates were compared against the progenitor strain (PAO1) to determine mutations selected following treatment with 8 μg mL^−1^ colistin. Called variants were visually inspected by mapping the reads on the reference genome (accession number NC_002516) using the software Artemis (http://sanger-pathogens.github.io/Artemis/Artemis/).

### Cloning and complementation

The wild-type *wzy* ORF was PCR-amplified as outlined in *supplementary information 7* from wild-type PAO1 genomic DNA template. The PCR product was purified using a GeneJET PCR Purification Kit (Thermo Scientific™) and cloned into plasmid pUCP20 plasmid at the restriction sites SacI and SphI. *E. coli* DH5α electrocompetent cells were prepared and electroporated following the protocol reported of Dower *et al* (Dower et al., 1988). Positive transformants were confirmed by PCR and the plasmid was extracted using a GeneJET Plasmid Miniprep Kit (Thermo Scientific™). Electrocompetent cells of *Pseudomonas aeruginosa* (PAO1, isolate D, and isolate F) were prepared following the protocol described by Choi *et al* (Choi et al., 2006). Complementation of the colistin-resistant phenotype was confirmed by measuring the minimal inhibitory concentration (MIC) of colistin against PAO1, and isolates D and F alone, and against these strains containing the empty vector or containing pUCP20-*wzy*. After incubation overnight, 10 μL of 0.02% resazurin was added to each well of the MIC plates to detect cell viability and determine the minimal inhibitory concentration.

### Statistical analysis

Unless otherwise stated, all data represent the mean ± SD of three independent biological experiments. Statistical differences in CFU mL^−1^ counts between single-species cultures were analysed by one-way ANOVA and differences in CFU mL^−1^ counts between microbial co-cultures were analysed by two-way ANOVA using GraphPad Prism version 8.2.0. Statistical analysis for the real-time PCR experiments was performed in R V2.0. To determine whether the data was normally distributed, Shapiro-Wilk tests were performed taking as input the expression values for each gene in each time point in the two different conditions (continuous-flow and batch culture). As the tests showed that data were normally distributed, Bartlett’s tests were used to assess whether the samples had the same variance. As the latter showed heterogenous variances, two-sided Welch’s T-test was used to calculate p-values of gene expression in continuous-flow vs batch culture conditions. P-values lower than 0.05 were considered statistically significant.

## Data availability statement

The raw data supporting the conclusions of this manuscript will be made available by the authors, without undue reservation, to any qualified researcher. The Illumina reads of the colistin-resistant isolates and parental strain were deposited in GenBank under accessions SAMN18614912 to SAMN18614920.

## Author contributions

TJO’B conceived and designed the work, executed the experiments, analysed the data, performed statistical tests and drafted/revised the manuscript. WF performed the WGS and associated genomics data analysis, performed statistical tests, cloned *wzy* and executed complementation experiments, determined MICs of the complemented strains, and contributed to drafting the manuscript. MW secured funding for the work, analysed the data, and revised the manuscript.

