## Supplementary information for "The efficacy of antimicrobial agents is decreased in a polymicrobial environment"

#### Supplementary Information 1

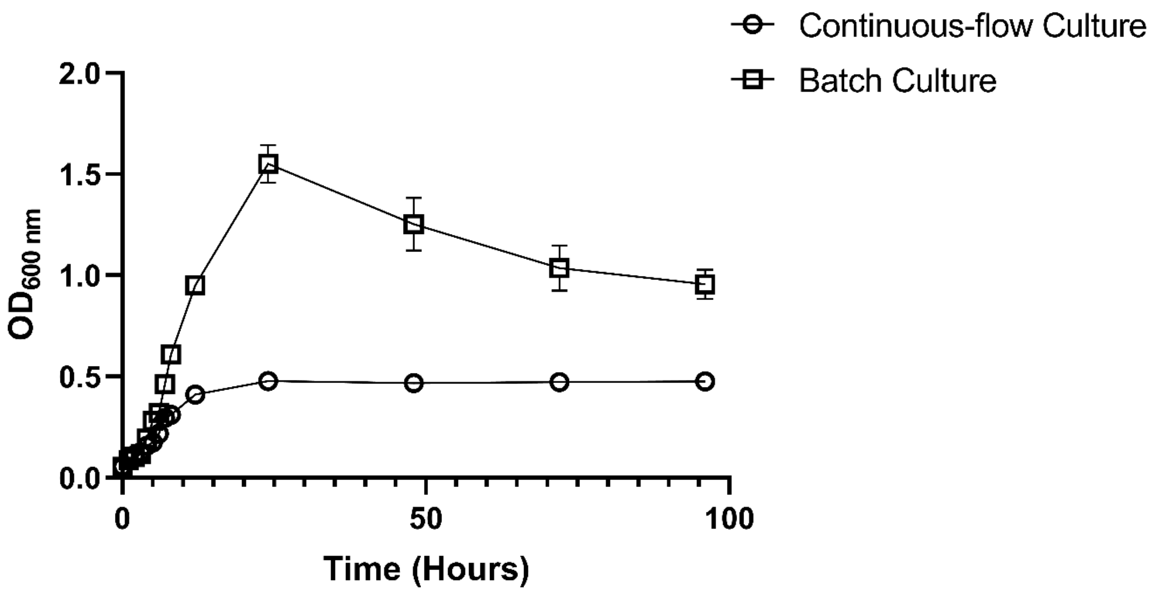


**Figure S1 Growth of *P. aeruginosa* in ASM.**

Growth of *P. aeruginosa* PAO1 in ASM under both batch and continuous-flow culture conditions. Growth is monitored by optical density ($OD_{600\ nm}$) during **(A)** continuous-flow culture ($Q$ = 170 µL $min^{-1}$); **(B)** batch culture ($Q$ = 0 µL $min^{-1}$). Data represent the mean ± standard deviation from three independent experiments.

#### Supplementary Information 2

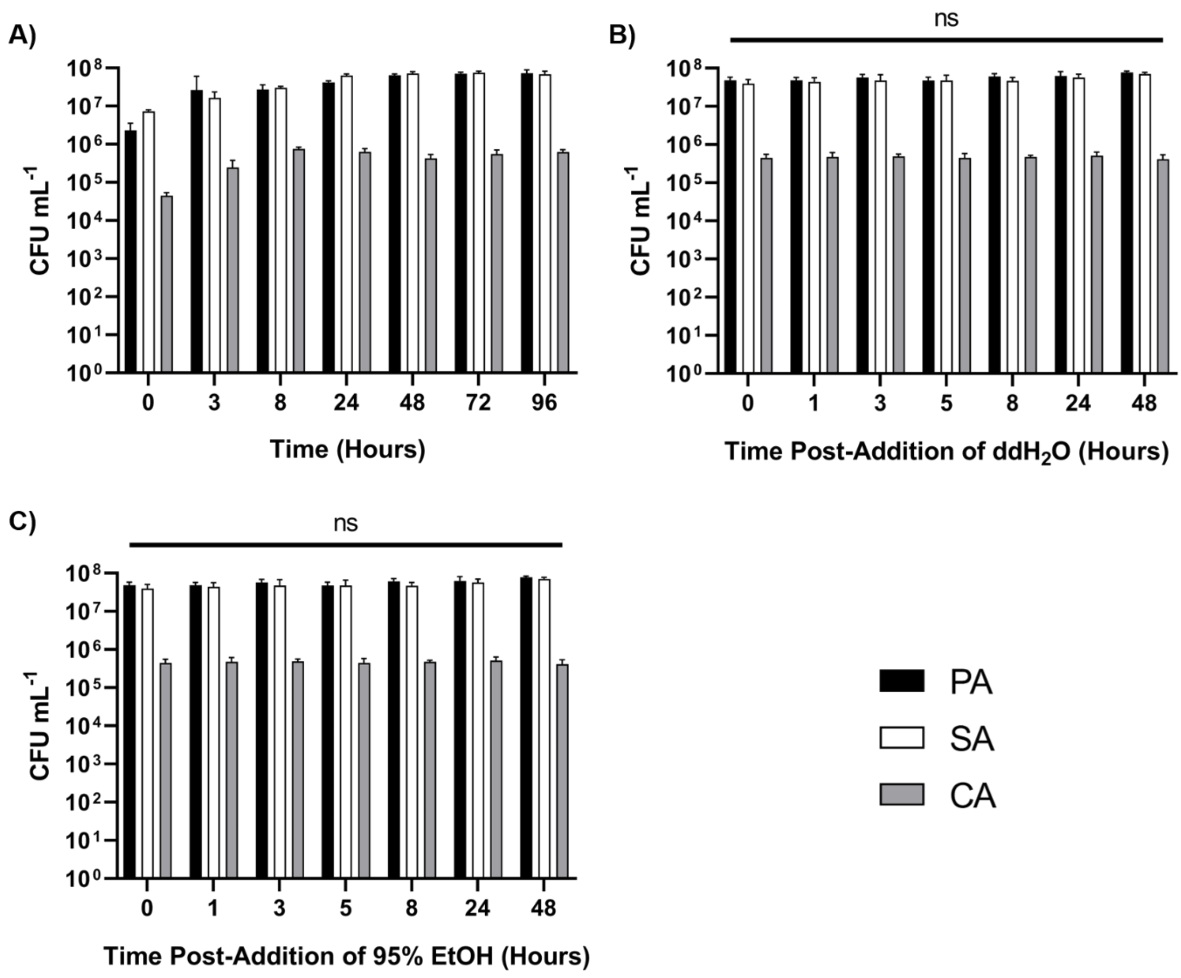


**Figure S2. Establishment of steady-state microbial communities and their treatment with antimicrobial solvents.** Viable cell counts of *P. aeruginosa* PAO1 (PA, black bars) *S. aureus* 25923 (SA, white bars) and *C. albicans* SC5314 (CA, grey bars) co-cultured in ASM under continuous-flow conditions. Flow rate = 145 µL min$^{-1}$. **(A)** Polymicrobial cultures reach a steady-state by T = 24 h, following this there is no significant difference in the CFU mL$^{-1}$ counts of any species. Established steady-state polymicrobial cultures (incubated for 24 h) were then treated with **(B)** 1 mL water or **(C)** 1 mL 95% ethanol used to dissolve colistin or fusidic acid/fluconazole, respectively. There was no significant change in CFU mL$^{-1}$ of any species following the addition of the two solvents (at T = 0 h). Data represented as mean ± standard deviation of three independent experiments. CFU mL$^{-1}$ values are plotted on a $log_{10}$ scale and $P$ values > 0.05 are considered as not significant (ns).

### Supplementary Information 3

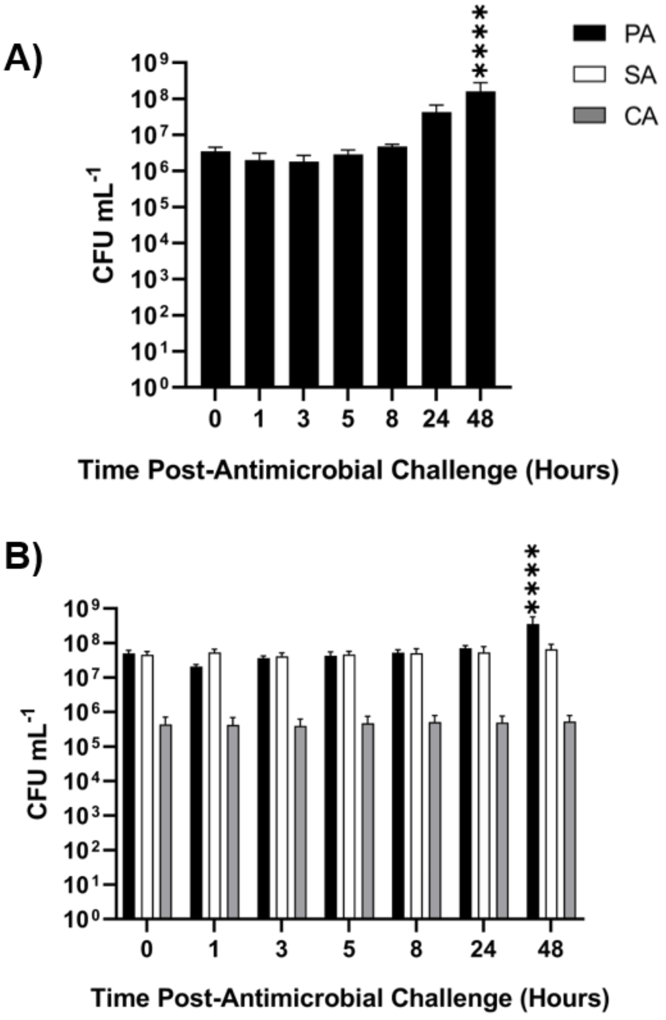


**Supplementary information 3 Addition of 1 x MIC colistin to steady-state microbial cultures.**

Mono-species *P. aeruginosa* PAO1 (PA) and mixed-species cultures of *P. aeruginosa* PAO1 (black bars), *S. aureus* 25923 (white bars) and *C. albicans* SC5314 (grey bars) were grown to a steady-state in ASM under continuous-flow conditions. Bars represent viable cell counts (CFU $mL^{-1}$) within **(A)** mono-species PA and **(B)** mixed-species populations following the addition of 4 µg $mL^{-1}$ colistin to the culture vessel. Data represented as the mean ± standard deviation from three independent experiments. Asterisks represent significant (**** $P < 0.0001$) differences in PA CFU $mL^{-1}$ counts in comparison to counts at the 0 hrs timepoint.

### Supplementary Information 4

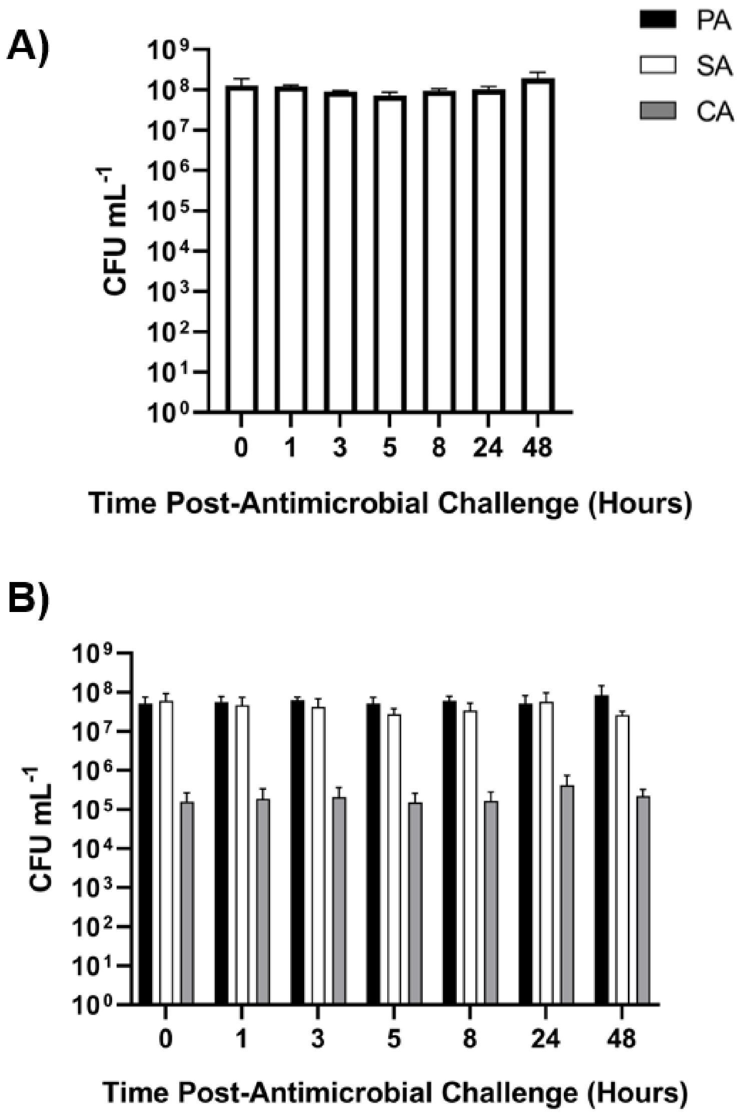


**Supplementary information 4 Addition of 1 x MIC fusidic acid to steady-state microbial cultures.**

Mono-species *S. aureus* 25923 (SA) and mixed-species cultures of *P. aeruginosa* PAO1 (black bars), *S. aureus* 25923 (white bars) and *C. albicans* SC5314 (grey bars) were grown to a steady-state in ASM under continuous-flow conditions. Bars represent viable cell counts (CFU $mL^{-1}$) in steady-state (24 hrs) **(A)** mono-species SA and **(B)** mixed-species populations following the addition of 15.6 ng $mL^{-1}$ fusidic acid to the culture vessel at T = 0 hrs. Data represented as the mean ± standard deviation from three independent experiments.

### Supplementary Information 5

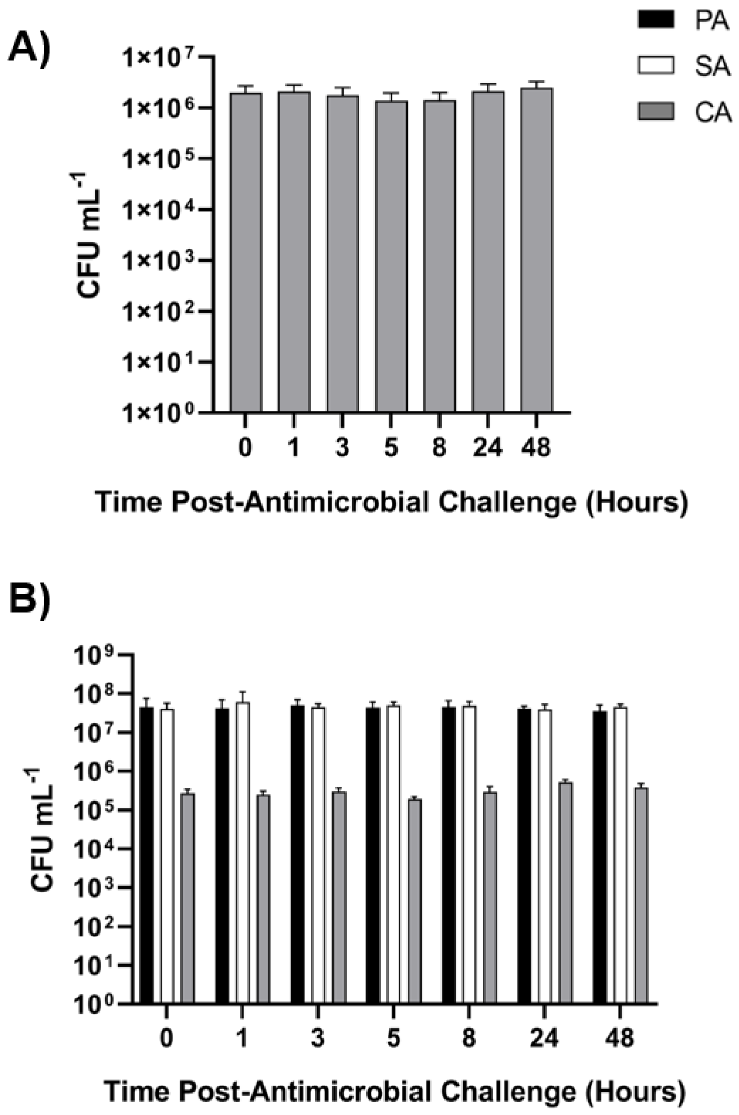


**Supplementary Information 5 Addition of 1 x MIC fluconazole to steady-state microbial cultures.**

Mono-species *C. albicans* SC5314 (CA) and mixed-species cultures of *P. aeruginosa* PAO1 (black bars), *S. aureus* 25923 (white bars) and *C. albicans* SC5314 (grey bars) were grown to a steady-state in ASM under continuous-flow conditions. Bars represent viable cell counts (CFU $mL^{-1}$) in steady-state (24 hrs) **(A)** mono-species CA and **(B)** mixed-species populations following the addition of 1 µg $mL^{-1}$ fluconazole to the culture vessel at T = 0 hrs. Data represented as the mean ± standard deviation from three independent experiments.

### Supplementary Information 6 Quantitative Real-time PCR (RT-PCR)

To determine the metabolic state and growth phase of steady-state populations maintained under continuous flow conditions, the relative expression of four stationary phase-specific and four exponential-phase specific *Pseudomonas aeruginosa* genes were selected for RT-PCR from a previously published transcriptomic dataset (Mikkelsen et al., 2007). By targeting eight genes encoding different products differentially expressed between the exponential and stationary phases of growth, discrepancies in the expression of genes due to differences in growth media (artificial sputum media vs AGSY) should be mitigated. Thus, making general trends regarding the metabolic state of the culture obvious. All RT-PCR results were normalised by quantifying the expression of the constitutively expressed 16S rRNA gene encoding for the RNA component of the 30S ribosomal subunit (Clarridge, 2004, Kolbert and Persing, 1999, Woese, 1987) and results analysed using the comparative $\Delta\Delta C_t$ method as described by (Giulietti et al., 2001). *Supplementary Information 6.1* shows the target genes, predicted gene products and *Supplementary Information 6.2* shows the reaction mixture and thermocycler conditions used for RT-PCR amplification.

#### RT-PCR primer design

Primer-BLAST software (NCBI, www.ncbi.nlm.nih.gov/tools/primer-blast/) was used to identify suitable primer pairs against the target genes with an approximate $T_m$ of 58°C and yielding an amplicon product of 150 bp.

#### RT-PCR reactions

RT-PCR amplification was carried out in MicroAmp Optical 96-well Reaction Plates (Applied Biosystems) sealed with MicroAmp Optical Adhesive Film (Applied Biosystems) using a 7300 Real-Time PCR System (Applied Biosystems). Reactions were carried out in 20 µL total volumes, using Universal PowerUp SYBR Green Master Mix (Applied Biosystems), following the manufacturer's instructions and using ROX as a passive reference dye. The reaction mixture and cycle conditions used for all RT-PCR reactions is displayed in *Supplementary Table 4*. RT-PCR of the 16S rRNA housekeeping gene was performed for each cDNA sample tested on every 96-well reaction plate to normalise results (and account for plate-to-plate variability between reactions. Amplicon products were resolved on a 1.4% (w/v) agarose gel to check that a single band of approximately 150 bp in size was present (data not shown).

| Primer Name | Target Gene | Gene Product | 5' - 3' Sequence | Growth Phase |
|---|---|---|---|---|
| rpsM-F | *rpsM* | 30S ribosomal subunit protein S13 | CGTCGCGAAATCAACATGAAC | *E* |
| rpsM-R | | | TTACTTGCGGATCGGCTTAC | |
| rplM-F | *rplM* | Early assembly protein of the 50S ribosomal subunit | TACCACCACTCCGGCTTC | *E* |
| rplM-R | | | CACCTTCAGCTTGCGATACA | |
| rpoA-F | *rpoA* | DNA-directed RNA polymerase α-chain | GCACCGAAGTGGAACTGTTG | *E* |
| rpoA-R | | | CAGTGGCCTTGTCGTCTTTC | |
| sodB-F | *sodB* | Superoxide dismutase | AACACCTACGTGGTGAACCT | *E* |
| sodB-R | | | GCTCAGGCAGTTCCAGTAGA | |
| rmf-F | *rmf* | Ribosomal modulation factor | ACGGCATAACCGGTAAATCTC | *S* |
| rmf-R | | | GCTGGAGTTGATTGAGACGTT | |
| rsmA-F | *rsmA* | Putative carbon storage regulator | CCCTGATGGTAGGTGACGAC | *S* |
| rmsA-R | | | GGTTTGGCTCTTGATCTTTCTC T | |
| rpoS-F | *rpoS* | Alternative sigma factor | AAGCTCGACCACGAACCTT | *S* |
| rpoS-R | | | CGTATCCAGCAGGGTCTTGT | |
| sodM-F | *sodM* | Superoxide dismutase | CTTCGAGGCGTTCAAGGATG | *S* |
| sodM-R | | | ATCGGCGTATTGCCGTTC | |
| F-16SrRNA-Pa-RT | 16S rRNA | RNA component of 30S ribosomal subunit | ACACTGGAACTGAGACACG | *C* |
| R-16SrRNA-Pa-RT | | | AGACCTTCTTCACACACG | |

**Supplementary Information 6.1 Target genes and primer pairs designed for RT-PCR analysis of the metabolic state of steady-state microbial cultures.**

Forward and reverse primers for a specific target gene are denoted by '-F' or '-R' respectively and growth phase indicates at what point within a typical growth curves these genes are expressed: (E) exponential phase, (S) stationary phase and (C) constitutively expressed.

| Reaction mixture | | |
|---|---|---|
| **Component** | **Volume** | **Final Concentration** |
| 2 x PowerUp SYBR Green Master mix | 10 µL | |
| Forward Primer | 1 µL | 0.25 µM |
| Reverse Primer | 1 µL | 0.25 µM |
| cDNA Template | 0.4 µL | 10 ng µL$^{-1}$ |
| Nuclease-free water | 7.6 µL | |
| **Thermocycler conditions** | | |
| **Temperature (°C)** | **Time** | **No. Cycles** |
| 50 | 2 min | Hold |
| 95 | 2 min | Hold |
| 95 | 15 secs | 40 |
| 60 | 2 min | |
| 4 | ∞ | Hold |

**Supplementary Information 6.2 RT-PCR reaction mixture and conditions.**

Reaction mixture (20 µL total volume) and thermocycling conditions for RT-PCR of target genes using universal PowerUP SYBR Green Master Mix kit and a 7300 Real-Time PCR System.

### Supplementary Information 7

| **Primer Name** | **Target Gene** | **Gene Product** | **5' - 3' Sequence** |
|---|---|---|---|
| wzy-F | *wzy* | B-band O-antigen polymerase | ATCCGG *GAGCTC*<br>AGGAGGAACAGCA**ATG**TATAT<br>ACTTGCTCGAGTCGACA |
| wzy-R | | | ATCCGG *GCATGC*<br>**TCA**TAGAGTTTTTCCTAAAGAC<br>ATCTTGA |

**Supplementary Information 7.1 Oligonucleotides used for cloning *wzy* in pUCP20.**

Sequences of forwards and reverse primers to amplify and clone *wzy* in pUCP20. Restrictions sites SacI and SphI are shown in italics, underlined sequences correspond to the ribosome binding site (RBS), and letters in bold represent the start and stop codons.

| **Reaction mixture** | | |
|---|---|---|
| **Component** | **Volume** | **Final Concentration** |
| 5X Q5 Reaction Buffer | 10 µL | 1x |
| 5X Q5 High GC Enhancer | 5 µL | 0.5x |
| Forward Primer | 1.25 µL | 0.25 µM |
| Reverse Primer | 1.25 µL | 0.25 µM |
| dNTPs | 1 µL | 0.2 mM |
| Q5 High-Fidelity Polymerase | 0.5 µL | 0.02 U/µl |
| Template DNA | 40 ng | 0.8 ng $\mu L^{-1}$ |
| Nuclease-free water | To 50 µL | |

| Thermocycler conditions | | |
|---|---|---|
| **Temperature (°C)** | **Time** | **No. Cycles** |
| 95 | 3 min | 1 |
| 95 | 15 min | 35 |
| 57 | 15 secs | |
| 72 | 1 min | |
| 4 | ∞ | Hold |

**Supplementary Information 7.2 PCR reaction mixture and conditions.**

Reaction mixture (50 µL total volume) and thermocycling conditions for PCR amplification of *wzy* for cloning in pUCP20 using Q5 high-fidelity polymerase.
